# Behavioral and neural decomposition of skull-induced death awareness

**DOI:** 10.1101/2020.03.31.018309

**Authors:** Tianyu Gao, Yue Pu, Jingyi Zhou, Guo Zheng, Yuqing Zhou, Shihui Han

## Abstract

Death awareness influences multiple aspects of human lives, but its psychological constructs and underlying brain mechanisms remain unclear. We address these by measuring behavioral and brain responses to images of human skulls. We show that skulls relative to control stimuli delay responses to life-related words but speed responses to death-related words. Skulls compared to the control stimuli induce early deactivations in the posterior ventral temporal cortex followed by activations in the posterior and anterior ventral temporal cortices. The early and late neural modulations by perceived skulls respectively predict skull-induced changes of behavioral responses to life- and death-related words and the early neural modulation further predicts death anxiety. Our findings decompose skull-induced death awareness into two-stage neural processes of a lifeless state of a former life.

**One sentence summary:** Behavioral and brain imaging findings decompose skull-induced death awareness into two-stage neural processes of a lifeless state of a former life.

## Main

Death awareness (DA) is a unique mental activity that emerges during human childhood^1^ and affects multiple aspects of our lives including art, religion, and medical care^2-4^. DA not only provides a framework for philosophical thinking about life^5^ but also influences human behavior, mental health^6-10^, and relevant brain activities^11-14^. Distinct schools of thought even regard DA as one of the defining traits of Homo sapiens^15^. Despite the broad impact of DA, its psychological constructs and underlying brain mechanisms have been poorly understood.

Conceptually, death anxiety and death reflection are supposed to represent two distinct forms of DA^16^. Empirically, however, it is difficult to clarify psychological constructs and neural processes involved in DA because it is capricious regarding when and how it occurs to mind. Behavioral research sought to manipulate DA by asking individuals to complete words from letter strings, to write sentences about one’s own death, or to watch videos of deadly automobile accidents^16^. Brain imaging studies revealed increased frontal/parietal and amygdala activity^17-20^ but decreased cingulate/insular activities^12,17, 19,21^ in responses to death-related words or sentences. These findings, however, did not disentangle neurocognitive processes underlying DA from upstream semantic processing and downstream cognitive/affective effects associated with the language stimuli.

Here we investigated neurocognitive underpinnings of DA by measuring behavioral and brain responses to images of human skulls. Skulls were treated in a special status in human history^22,23^. Images of skulls provide reliable visual cues for death detection^24^ and have been used to remind people of life’s finiteness^25^. These facts reflect a pervasive belief that skulls may bring *death* into consciousness and suggest images of skulls as good candidates for investigation of neurocognitive processes involved in DA. A skull reminds us of dissolution of the soul-body complex, which, according to some philosophical thoughts^26^, manifests the essence of death. A skull is different from both animate (e.g., faces) and inanimate (e.g., house) stimuli used in previous research^27^ in that it was *animate* before but is *inanimate* now. Therefore, a dualistic mental experience of a lifeless (or an inanimate) state of a former life (or an animate entity) may reflect the nature of skull-induced DA.

### Behavioral decomposition of DA

We tested this hypothesis in Experiment 1 by assessing how images of skulls affect reaction times (RTs) to classification of death-related and life-related words in healthy adults (N=38, see Table S3 and SI for selection of word stimuli). We predicted that skull-induced DA would produce opposite effects on the processing of the two types of words by speeding responses to death-related words but slowing responses to life-related words due to conceptual congruency regarding death or incongruency regarding life between skulls and the words. On each trial of the classification task, a black-and-white drawing of a human skull preceded each word as a prime (Figure 1A). The primes also included neutral and fearful faces to control for possible effects of face-like configural structure and negative affect produced by skulls. Inverted skulls and scrambled stimuli derived from drawings of skulls and faces were used as primes to control for possible effects of perceptual feature processing and intrinsic differences in RTs to death-related and life-related words.

**Figure 1.**
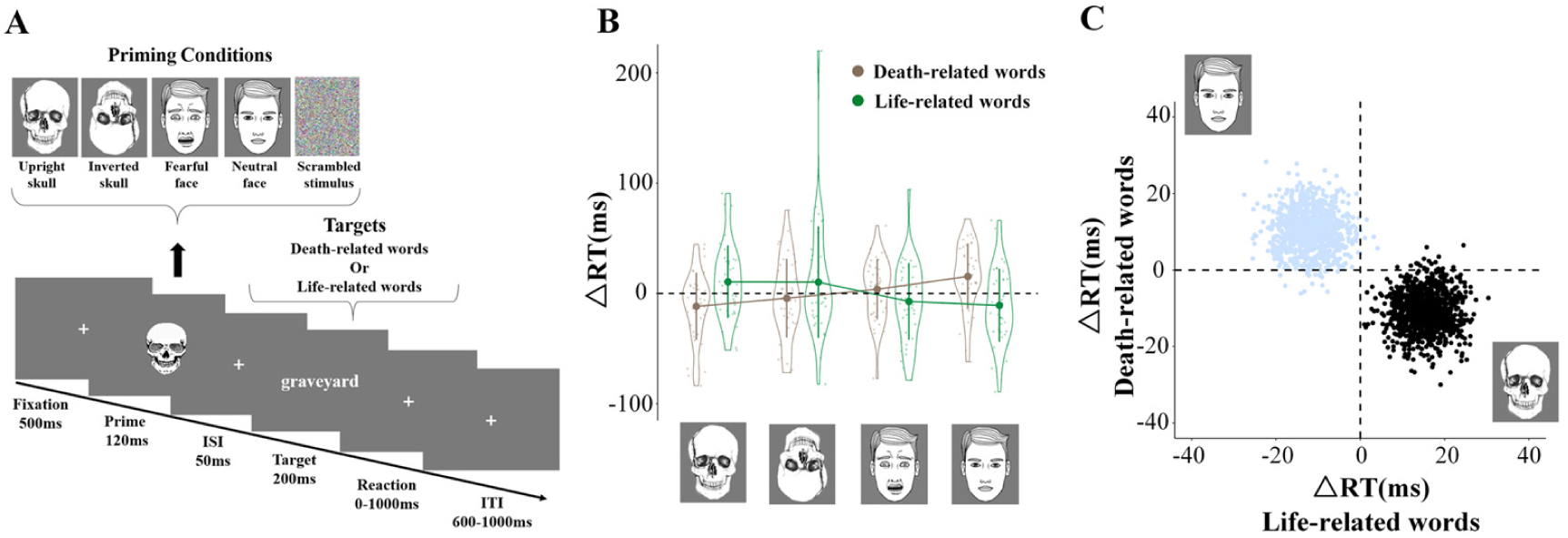
Experimental procedures and results in Experiment 1. (A) Stimuli and procedure in each trial. Five types of primes are illustrated. (B) ΔRTs to death- and life-related words preceded by skulls and faces (vs. scrambled stimuli). Shown are group means (big dots), standard deviation (bars), measures of each individual (small dots), and distributions (violin shape) of ΔRTs. (C) The results of a bootstrap analysis. This analysis calculated a new data set using nonparametrically resampling with replacement from the data set of ΔRTs to death-related and life-related words in the skull and neutral face priming conditions. The mean of this bootstrapped sample was plotted as a point in a 2-D space with x and y axes showing ΔRTs to death-related and life-related words, respectively. The bootstrapped sample mean points (based on 1,000 repetitions) in the skull and face conditions fall separately in the 2-D plot.

To examine the priming effect in each condition, we calculated differential RTs (ΔRTs) by subtracting RTs in the scrambled prime condition from other priming conditions. A repeated measure analysis (ANOVA) of ΔRTs with Prime (upright skull, inverted skull, fearful face, neutral face) and Word type (death-related vs. life-related) as within-subjects factors showed a significant two-way interaction (F(3,111) = 17.02, p < 0.0001, η_p_^2^ = 0.315, 90% CI = [0.187, 0.404], Figure 1B, Table S4). Post-hoc comparisons revealed that upright skulls tended to speed responses to death-related words but delayed responses to life-related words, resulting in a significant difference in RTs between death- and life-related words (Mean difference = −22.25, Bonferroni corrected p = 0.008, 95% CI = [−6.22, −38.28]). Neutral faces, however, produced significantly opposite effects on RTs to death- and life-related words (Mean difference = 26.35, p < 0.001, 95% CI = [13.71, 38.98]), whereas no significant priming effect was observed for inverted skulls and fearful faces (Mean difference = −14.51 and 11.10, p = 0.115 and 0.109, 95% CI = [−32.71, 3.69] and [−2.59, 24.78]). The opposite priming effects of skulls/neutral faces on death-related and life-related words were further highlighted using a bootstrap analysis^28^ (see Figure 1C), and were replicated in the following neuroimaging experiments (Figure S4). However, images of inanimate objects such as houses failed to produce opposite priming effects on death-related and life-related words (see SI results of Experiment 2, Figure S5).

The priming effects on categorization are believed to reflect that recently activated psychological constructs are retrieved and used to process a subsequent stimulus^29,30^. Our behavioral findings have two important implications. First, skulls and faces activate different psychological constructs that influence following semantic categorization of death-related and life-related words in opposite directions. Second, skull-activated psychological constructs can be used during subsequent processes to facilitate classification of death-related words but to inhibit classification of life-related words regardless the difference in perceptual processing of images of skulls and words. These findings provide evidence that images of skulls activate mental experiences that link to both death and life and support our hypothesis that skull-induced DA essentially reflects a dualistic mental process of a lifeless state of a former life.

### Two-stage neural modulations by perceived skulls

Next we examined brain activity underlying skull-induced DA in Experiment 3 by recording electroencephalography (EEG) from an independent sample (N=30) who viewed rapid presentations of black-and-white drawings of skulls and neutral faces in one session and of fearful and neutral faces in another session (see SI Methods). Inverted skulls and faces were used to control for effects of perceptual features of each stimulus set. On each trial a stimulus was displayed for 200 ms and followed by a fixation cross (duration = 800 ∼ 1200 ms). The participants performed a one-back task by pressing a button to a causal repetition of the same stimulus in two consecutive trials (10% of all trials).

Event-related potentials (ERPs) with a millisecond resolution were obtained by averaging EEG to stimuli of each category. The results showed that upright compared to inverted skulls elicited a negative-going activity at 120-140 ms post-stimulus at the occipital/parietal region (F(1,29) = 107.212, p < 0.0001, η^2^_p_ = 0.787, 90% CI = [0.647, 0.846], Figure 2A). We called this activity death-awareness related negativity (DRN). The DRN amplitude significantly distinguished between skulls and neutral faces (F(1,29) = 77.088, p < 0.0001, η^2^_p_ = .727, 90% CI = [0.556, 0.803]) despite their similarity in configural structures. Neural responses to upright (vs. inverted) fearful faces were characterized by two negativities at 104-124 ms (Nd1 at the occipital/parietal region) and 188-208 ms (Nd2 at the central/frontal region) (F(1,29) = 22.423 and 37.386, ps < 0.0001, η^2^_p_ = 0.436 and 0.563, 90% CI = [0.200, 0.587] and [0.337, 0.683]). The Nd1 and Nd2 amplitudes differed significantly between fearful and neutral faces (F(1,29) = 25.088 and 26.53, ps < 0.0001, η^2^_p_ = 0.464 and 0.478, 90% CI = [0.226, 0.608] and [0.240, 0.619]). The DRN results provide evidence for an early neural modulation that characterizes the process of skulls.

**Figure 2.**
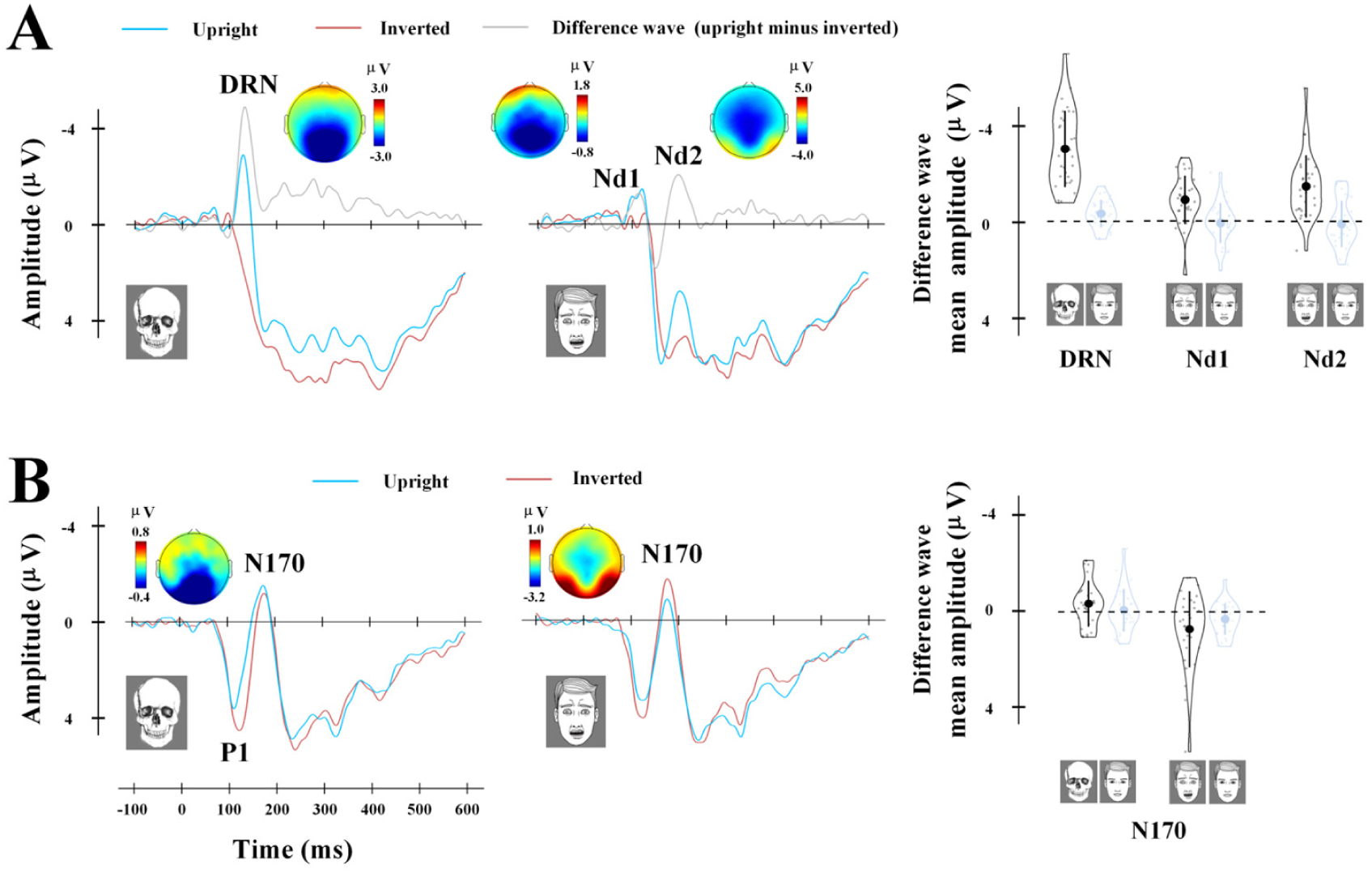
EEG results in Experiment 3. (A) ERPs to skulls and fearful faces at the middle occipitoparietal electrodes. (B) ERPs to skulls and fearful faces at the right occipitoparietal electrode. The voltage topographies show scalp distributions of the difference wave (upright minus inverted skulls or fearful faces) in DRN/Nd1/Nd2/N170 time windows. Group means (big dots), standard deviation (bars), measures of each individual (small dots), and distributions (violin shape) of the difference waves or N170 amplitudes are shown in the right panels.

Both skulls and fearful faces elicited a negative activity over the bilateral occipital/temporal regions at 160-200 ms post-stimulus. This activity, designated as N170, is usually larger to inverted than upright faces and this face-inversion effect reflects recruitment of additional non-face mechanisms to process inverted faces^31^ (Yovel, 2016). We assessed the inversion effect on the N170 amplitudes to faces and skulls by conducting ANOVAs of the mean N170 amplitudes with EEG session (skull/neutral-faces vs. fearful/neutral-faces), Stimuli (skull/fearful vs. neutral faces) and Orientation (upright vs. inverted) as within-subjects variables. The results showed a significant three-way interaction (F(1,29) = 8.29, p = 0.007, η^2^_p_ = 0.222, 90% CI = [0.037, 0.407]). Post-hoc comparisons revealed a significantly larger N170 amplitude to inverted (vs. upright) fearful faces (Mean difference = −0.832 μV, p = 0.006, 95% CI = [−1.41, −0.26]) but a significant enhancement of the N170 amplitude to upright (vs. inverted) skulls (Mean difference = 0.436 μV, p = 0.015, 95% CI = [0.09, 0.78], Figure 2B). The N170 enhancement to upright (vs. inverted) skulls revealed a second-stage neural modulation that distinguished skulls from faces.

### Replication of two-stage neural modulations by perceived skulls

Because skulls and fearful faces were presented in different sessions in Experiment 3, the contextual effects produced by neutral faces might be different for brain activities in response to skulls and fearful faces. In Experiment 4 we compared ERPs to skulls, fearful and neutral faces which were presented in the same EEG session in an independent sample (N=28). In Experiment 5 we further tested whether perception of any inanimate objects would produce similar two-stage neural modulations produced by images of skulls by recording EEG to black-and-white drawings of skulls, faces, and houses from an independent sample (N=36). We also examined whether perceived images of real human skulls produce similar two-stage neural modulations in Experiment 6 by recording EEG from an independent sample (N = 30). The results in Experiments 4-6 replicated the two-stage neural modulations by skulls (see SI for methods and results, Figure 3A, Figures S5 and S6). However, the ERPs to houses did not show DRN and N170, indicating that not any inanimate object can generate the DRN.

**Figure 3.**
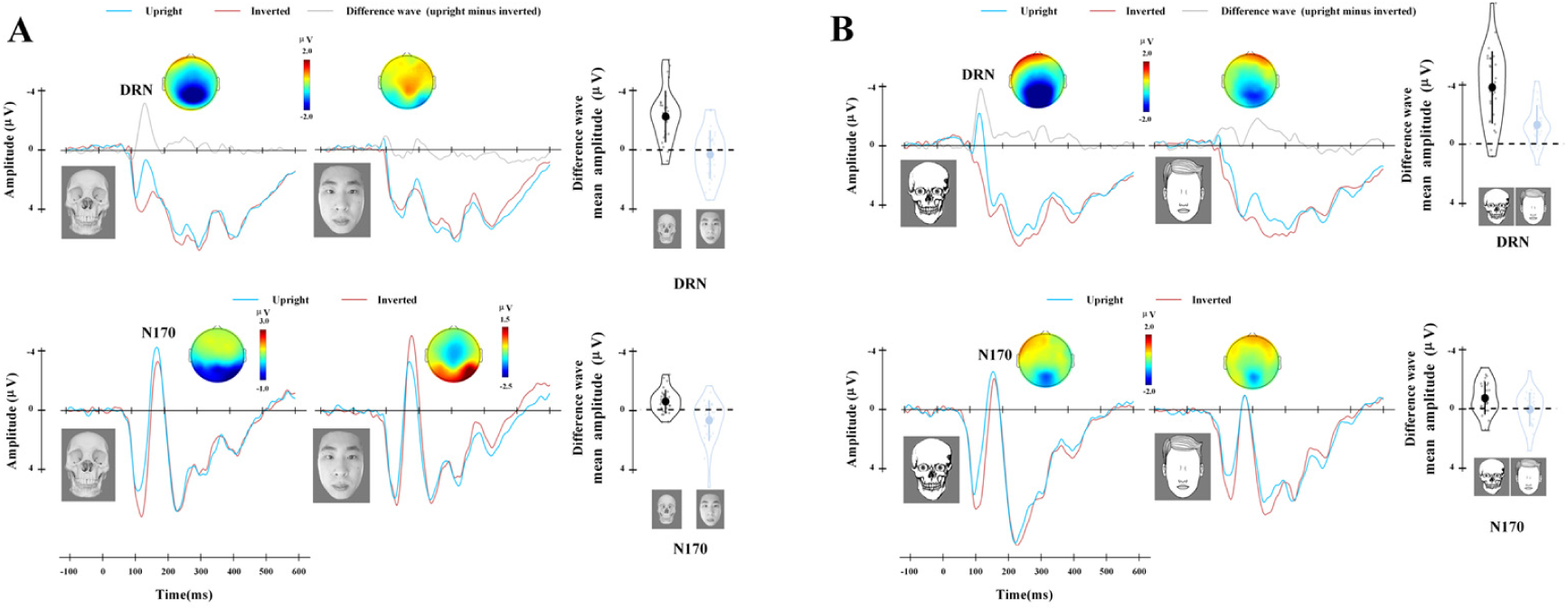
Results of Experiments 6 and 7. (A) The DRN and N170 to images of human skulls and faces. (B) The DRN and N170 to skulls with eyeballs and faces without eyeballs. The voltage topographies show scalp distributions of the difference wave (upright minus inverted skulls or faces) in DRN and N170 time windows. Group means (big dots), standard deviation (bars), measures of each individual (small dots), and distributions (violin shape) of the difference waves or N170 amplitudes are shown in the right panels.

Taken together, our EEG findings provide evidence for skull-specific two-stage neural modulations that are independent of contextual information and perceptual features of stimuli. The DRN differentiates skulls from inanimate objects (e.g., houses) and animate stimuli with similar configural structures (e.g., faces) and negative emotion (e.g., fearful faces). The late N170 also distinguishes skulls and faces by showing the opposite inversion effects on the N170 amplitudes to skulls and faces. The skull-induced neural modulations are also different from those related to artificial (vs. real) faces, which are characterized by a decreased lateral occipito-temporal activity (P1) at 100–140 ms post-stimulus^32^ (see SI for the skull-evoked P1 results in Experiment 3). To our knowledge, no previous research has reported the DRN and N170 enhancement effects that characterize the neural processing of perceived skulls. These findings revealed neural mechanisms underlying skull-induced mental experiences within 200 ms post-stimulus.

### Configural information of skulls and DRN

Because appearance of eyes is thought to be critical for assigning “life” to a face^33^, we further tested whether lack of eyeballs is pivotal for skull-induced two-stage neural modulations in Experiment 7. We recorded EEG from an independent sample (N = 28) while the participants performed the one-back task on black-and-white drawings of skulls with eyeballs and neutral faces without eyeballs (Figure 3B). If lack of eyeballs is sufficient and necessary for generation of the two-stage neural modulations, faces without eyeballs rather than skulls with eyeballs would produce DRN and N170 enhancement effects. The ERP results, however, showed a significant DRN (F(1,27) = 67.73, p < 0.001, η^2^_p_ = 0.715, 90% CI = [0.530, 0.796]) and N170 enhancement (F(1,27) = 10.85, p = 0.003, η^2^_p_ = 0.287, 90% CI = [0.069, 0.470]) to upright (vs. inverted) skulls with eyeballs (Figure 3B). Interestingly, upright vs. inverted faces without eyeballs elicited a significantly negative activity in the DRN window ((F(1,27) = 25.83, p < 0.001, η^2^_p_ = 0.489, 90% CI = [0.2426, 0.631], but this effect was significantly smaller than the DRN (F(1,27) = 24.05, p < 0.001, η^2^_p_ = 0.471, 90% CI = [0.224, 0.618]). Moreover, upright (vs. inverted) faces without eyeballs failed to increase the N170 amplitude (F(1,27) = 0.085, p = 0.772, η^2^_p_ = 0.003, 90% CI = [0.000, 0.097]), and the inversion effect on the N170 amplitude differed significantly between skulls with eyeballs and faces without eyeballs (F(1,27) = 7.92, p = 0.009, η^2^_p_ = 0.227, 90% CI = [0.0351, 0.417]). These results suggest that a local feature (e.g., lack of eyeballs) may contribute but not be sufficient to generate the two-stage neural modulations. Thus, configural information of a skull may play a fundamental role in generation of DA-related two-stage neural modulations.

### Sources of dynamic neural responses to skulls

Next we used magnetoencephalography (MEG) — a potential technique for millisecond source imaging^34^ — to estimate origins of the dynamic neural responses to skulls. Previous functional magnetic resonance imaging (fMRI) revealed distinct neural representations of animate and inanimate stimuli along the lateral to medial regions of the ventral temporal cortex (VTC)^27,35,36^ such as faces in the occipital face area (OFA)^37^ and fusiform face area (FFA)^38^, and houses in parahippocampal place area (PPA)^39^. Unlike the animate or inanimate stimuli used in previous research, a skull does not sit in either category alone but relate to both as a skull reminds a lifeless state of a former life. We predicted that images of skulls may evoke dynamic activities across multiple subregions in the VTC rather than limited to the lateral or medial part of the VTC that encodes animate or inanimate stimuli.

We tested this prediction in Experiment 8 by recording 306-channel, whole-head anatomically constrained MEG from an independent sample (N = 29). The stimuli and procedure were the same as in Experiment 3. Moreover, the participants completed the same behavioral test as in Experiment 1. Given our EEG findings, we first examined whether the temporal dynamics of sensor-space MEG signals that allows decoding of perceived skulls and faces. We conducted multivariate analyses using a linear support vector machine method for classification of upright vs. inverted skulls and upright vs. inverted fearful faces independently for each participant (see SI methods). The results showed that the decoding accuracy was significantly higher than the chance level (50%) for upright vs. inverted skulls at 80-400 ms (p < 0.001) and for upright vs. inverted fearful faces at 87-400 ms (p < 0.001, Figure 4A, Figure S8). Moreover, cluster-based t-tests confirmed a significantly higher decoding accuracy for upright vs. inverted skulls than upright vs. inverted fearful faces at 115-153 ms post-stimulus (p < 0.001). This is consistent with our finding of the DRN that distinguishes between skulls and faces in a similar time window.

**Figure 4.**
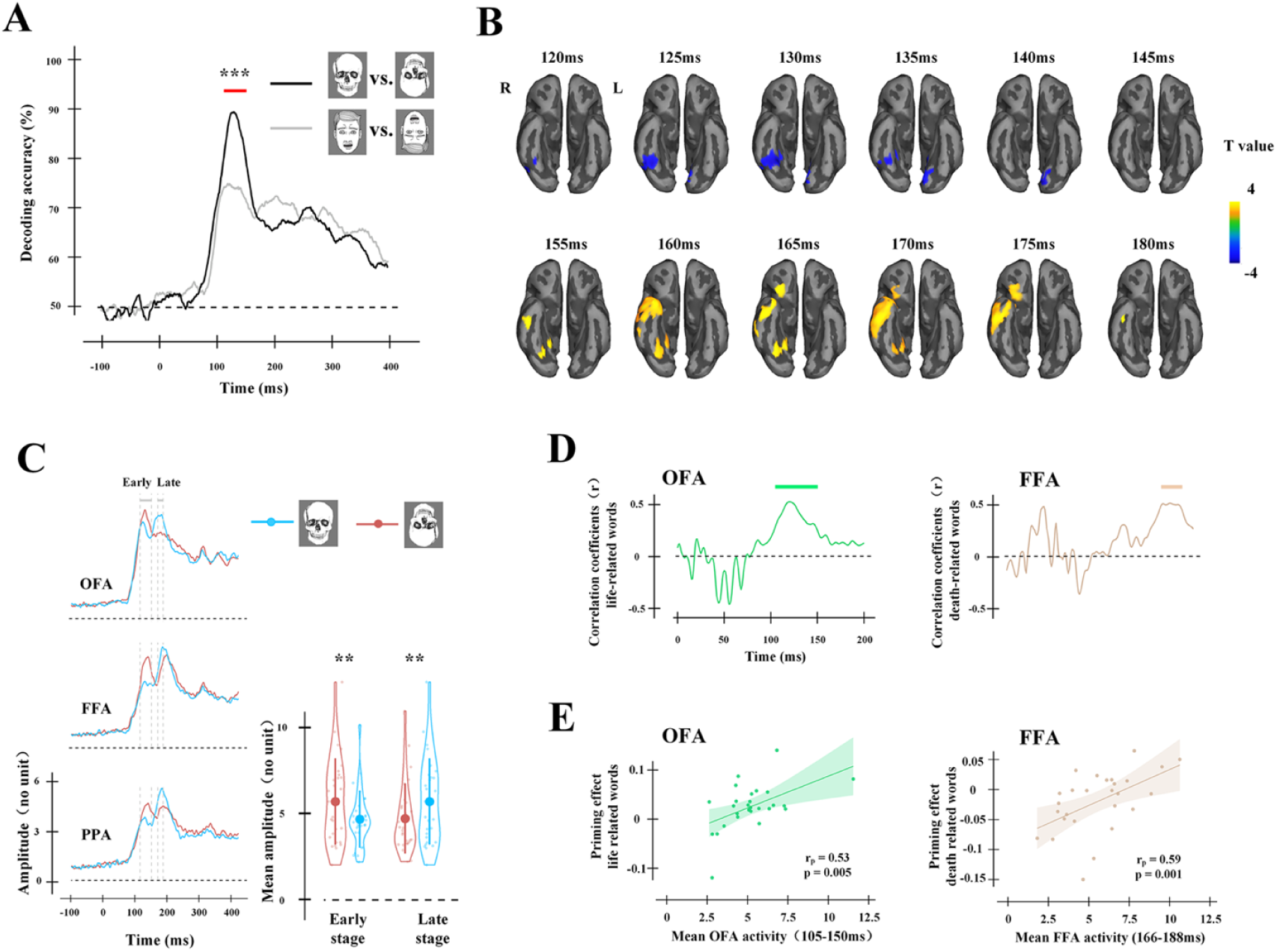
MEG results in Experiment 8. (A) Results of decoding accuracies. The red bar indicates time windows in which the decoding accuracy was significantly higher for upright vs. inverted skulls than upright vs. inverted fearful faces. (B) The neural origins of the two-stage modulations by upright (vs. inverted) skulls. These include the early decreased activity in the right posterior VTC and late increased activity in the right posterior VTC and ATL. Also see the supplementary video that illustrates the dynamic two-state neural modulations by skulls. (C) Source-space MEG signals in the independently fMRI defined OFA, FFA, and PPA. The left panel shows time courses of MEG signals to upright and inverted skulls. The right panel shows early decrease and late increase of the mean amplitudes of the MEG signals in these brain regions. (D) Results of point by point correlation analyses. The horizontal bars indicate time windows in which the OFA and FFA activity to uptight skulls predicted the priming effects on RTs to life-related and death-related words, respectively. (E) Correlations between the mean OFA/FFA activity and the priming effects on RTs to life-related/death-related words.

To localize the sources of neural modulations by skulls, we conducted a whole-brain analysis of MEG source-space signals to upright vs. inverted skulls at 100-200 ms based on our EEG results and MEG decoding results. The results revealed that, relative to inverted skulls, upright skulls induced significant deactivations in the right posterior VTC (110-140 ms, peak MNI coordinate x/y/z = 45/-57/-14, cluster-level corrected p = 0.024) and the left medial occipital area (120-145 ms, -5/-76/-8, cluster-level corrected p = 0.038, Figure 4B). This is followed by significantly increased activities in the right posterior VTC (153-173 ms, 23/-69/-10, cluster-level corrected p = 0.040) and right anterior temporal lobe (rATL) that extended into the right orbital frontal cortex (151-190 ms, 45/18/-16, cluster-level corrected p = 0.002). Similar whole-brain analyses showed significantly increased activities in the right posterior VTC (137-170 ms, 41/-67/-20; cluster-level corrected p = 0.002, Figure S9) to upright vs. inverted neutral faces but not fearful faces.

These MEG results uncover the origin of skull-induced neural modulations in the VTC, which is known to support neural representations of faces and house^27,38,39^. Our MEG results, however, did not provide direct evidence for associations of skull-induced DA with brain regions underlying representations of animate and inanimate stimuli such as the OFA, FFA, and PPA, which need to be functionally localized in each participant. Therefore, we scanned the participants using fMRI during perception of black-and-white drawings of faces and houses to localize these brain regions in each participant (see SI Methods). Similar to previous research^40^, the OFA and FFA were defined by contrasting faces vs. houses and the PPA was defined by the reverse contrast. To test potential associations of OFA/FFA/PPA activities with skull-induced DA, we extracted MEG source-space signals to upright and inverted skulls from these regions for each participant. Consistent with the results of the whole-brain analysis, the mean amplitudes of OFA/FFA/PPA source signals to skulls showed opposite patterns in the early (110-140 ms) and late (153-173 ms) time windows (F (2,54) = 24.77, p < 0.001, η^2^_p_ = 0.478, 90%CI = [0.300, 0.584], Figure 4C), due to early decreased activities to upright (vs. inverted) skulls (Mean difference = −0.906, p = 0.001, 95%CI=[−1.421, −0.390]) but late increased activities to upright (vs. inverted) skulls (Mean difference = 0.856, p = 0.006, 95%CI=[0.263, 1.449]). Such patterns did not differ significantly among the three regions (F (2,54) = 0.52, p = 0.595, η^2^_p_ = 0.019, 90%CI = [0, 0.029]). We then calculated point-by-point correlations between neural responses to upright skulls in each of these regions and ΔRTs to death-related and life-related words across individuals. The results revealed that greater OFA deactivation at 105-150 ms predicted faster responses to life-related words (all FDR corrected, Figure 4D and 4E), suggesting an association between early OFA deactivation and a mental experience of a former life elicited by perceived skulls. Moreover, greater FFA deactivations at 166-188 ms predicted faster responses to death-related words, suggesting an association between late FFA deactivations and a process of a lifeless state elicited by perceived skulls.

The MEG findings are consistent with our EEG results and further parsed the skull-specific two-stage neural modulations into early deactivations in the lateral and medial regions of the VTC in the DRN time window and late activations in the medial region of the VTC and rATL in the N170 time window. These results highlight the functional roles of OFA and FFA activities in skull-induced DA by showing evidence for associations between the OFA/FFA activity to skulls and skull-priming effects on behavioral responses to life-related and death-related words.

### Skull-induced brain activity and death anxiety

Finally, we tested whether skull-induced brain activity can predict subjective feelings of death-anxiety, which may reflect another form of DA^16^. We collected questionnaire measures of death anxiety (SI methods) from the participants in Experiments 3-5. Correlation analyses repeatedly revealed significant correlations between the DRN amplitudes to skulls and degree of death anxiety (Experiments 3-5: Pearson’s r = −0.384/-0.391/-0.381, p = 0.036/0.039/0.034, R^2^ = 0.1475/0.153/0.146, 90% CI = [0.0068, 0.3678] /[0.009, 0.374]/ [0.006, 0.366]), as individuals showing a larger skull-induced DRN reported greater death anxiety. These findings provide evidence for an association between early neural responses to skulls and general feelings and negative affect related to death.

## Discussion

Our findings demonstrate that images of skulls lead to two-stage neural modulations in the VTC, which overlaps with brain regions that mediate face perception (e.g., the OFA and FFA)^38,41^and represent multiple-level person knowledge (e.g., the ATL) ^42,43^. These neural modulations are different from neural responses to death-related words or sentences that activate the neural circuits underlying emotional responses (e.g., amygdala), emotion regulation (e.g. lateral frontal cortex and cingulate), and interoceptive representation (e.g., insula)^12,17-19,21^. These findings together indicate that skull-induced DA is essentially different from death-related thoughts induced by semantic stimuli in terms of their neural underpinnings. Our results further suggest that skull-related neural coding of a lifeless state of a former life in the posterior VTC may further activate semantic concept of death as a key component of person knowledge in the right ATL. The skull-induced neural modulations may interact with neural activities in other brain regions underlying cognitive and affective processes of life-related/death-related words and result in changes of behavioral performances during semantic categorization of the two types of words.

Perception of animate (e.g., faces) and inanimate (e.g., houses) stimuli is associated with activations the lateral and medial VTC, respectively^27^. Face perception also engages dynamic interactions between subregions of the VTC (e.g., OFA and FFA)^44^. Our findings revealed more complicated neural dynamics in response to skulls that activate a dualistic mental experience related to both animacy and inanimacy. Skull-elicited neural dynamics in the VTC demonstrate a new pattern of neural coding of a simple visual stimulus that engages both decreased and increased activities that spread across time and multiple regions in the VTC. This finding extends our understanding of the functional role of the VTC in supporting a dualistic mental experience of a lifeless state of a former life. Interestingly, the unique neural dynamics underlying perception of skulls was manifested mainly in the right hemisphere, suggesting a right hemisphere lateralization of neural activities supporting skull-induced DA. These findings cast new light on how the brain represents a visual stimulus that itself is inanimate but reminds us a former life.

Death anxiety is proposed to be one form of DA^16^, but there has been little evidence for association between death anxiety and brain activity underlying DA. Previous research suggests that decreased insular response to death-related words may covary with self-report of death anxiety but only in a small proportion of individuals who showed weak frontoparietal responses to death-related words^19^. Our findings indicate that death anxiety is associated with the early stage of neural modulations by perceived skulls. Therefore, subjective feelings of death anxiety may have neural origins in both the emotion processing system (e.g., insula) and cognitive processing system (e.g., the ventral VTC). These findings together open a new avenue to objective estimation of individual differences in DA-related emotional states.

Recent research of comparative thanatology has examined scent-triggered behavioral responses to dead conspecifics across different species of animals^15,45^. Central issues in this field include whether and how animals detect death in others and what cognitive processes contribute to the generation of DA. Since animals encounter skulls in the wild, our work suggests a new approach that does not rely on language stimuli but allows investigation of whether a similar neural system is evolved in non-human primates for detection of death. Addressing such questions helps to clarify the contention of DA as a defining trait of Homo sapiens^15^.

## Acknowledgments

We thank W. Li and S. Wang for work on stimulus figures.

## Author contribution

Conceptualization, S.H.; Methodology, S.H., T. G., Y. P, J. Z.; Investigation, T.G., Y.P., J.Z., G.Z., Y.Z., and S.H.; Resources, S.H.; Supervision, S.H.; Writing – original draft, S.H. and T.G.; Writing – review and editing, T.G., Y.P., J.Z., G.Z., Y.Z., and S.H..

## Code availability

The code used to analyze the data that support the findings of this study are available from the corresponding author upon reasonable request.

## Data availability

The data that support the findings of this study are available from the corresponding author upon reasonable request.

**Supplementary information** is available for this paper in the online version of the paper.

## Supplementary Information

### Pilot Experiment 1: Evaluation of black-and-white drawings of stimuli

Our study measured behavioral and neural responses to black-and-white drawings of human skulls to decompose skull-induced death awareness (DA). Black-and-white drawings of faces were used as controls. Black-and-white drawings were used in our work to remove detailed visual features of the stimuli so as to reduce the difference in these visual features between skulls and the control stimuli to a maximum degree.

To assess subjective feelings of emotion pertaining to black-and-white drawings of human skulls, we recruited 200 participants (84 males; mean age ± SD = 21.45 ± 3.17 yrs) in an online survey. Stimuli include 16 black-and-white drawings of human skulls from web photo galleries. All images were transformed to 300 x 360 pixels on a grey background (RGB:122, 122, 122), as illustrated in Figure S1. All participants in our study self-reported no neurological diagnoses, nonreligious, right-handed and had normal or corrected-to-normal vision. Written informed consent was obtained prior to the experiment. All participants were debriefed by explanation of the purpose of this study. This study was approved by the local ethics committee at the School of Psychological and Cognitive Sciences, Peking University.

**Figure S1.**
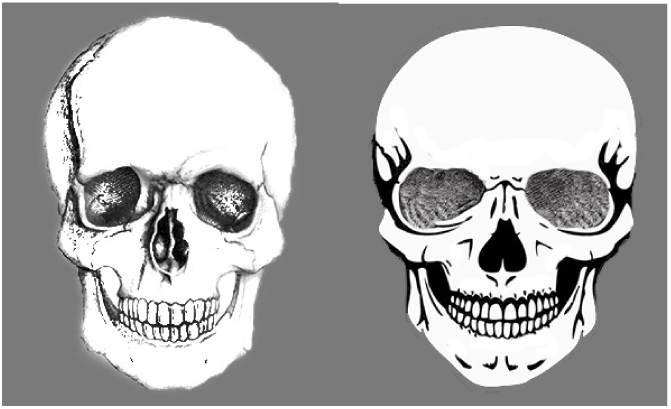
Illustrations of two black-and-white drawings of human skulls used in pilot Experiment 1.

Each participant was presented with 8 black-and-white drawings of human skulls and was asked to report whether a drawing evokes emotional responses such as happy, sad, angry, disgust, fear, or does not evoke a specific emotion (i.e., neutral). The participant was then asked to rate “how intense is your emotion evoked by the drawing?” (1 = very slight, 5 = moderate, 9 = extremely strong) and “how strongly are you aroused by the drawing?” (1 = very slightly, 5 = moderately, 9 = extremely strongly) on Likert scales. The participant was also asked to rate “How is the drawing related to death?” (1 = extremely unrelated, 5 = not sure, 9 = extremely related). Table S1 shows the frequencies of rating choices among the sample. The results suggest that black-and-white drawings of human skulls do not induce a specific dominant negative emotion. The emotional intensity and arousal related to these stimuli were slightly higher than moderate but were similar among the six types of emotions. Finally, explicit reports suggest weak subjective feelings of death-relatedness about perceived black-and-white drawings of human skulls.

**Table S1.** Rating results related to drawings of skulls in the pilot experiment

| Chosen Emotion | Frequency | Intensity<br>(Mean±SD) | Arousal<br>(Mean±SD) | Death-relatedness<br>(Mean±SD) |
| --- | --- | --- | --- | --- |
| Happy | 20.81% | 6.07(1.56) | 5.35(2.04) | 4.85(2.41) |
| Sad | 11.5% | 6.16(1.69) | 4.91(2.18) | 4.88(2.35) |
| Angry | 13.94% | 6.65(1.47) | 5.57(2.02) | 5.50(2.32) |
| Disgust | 12.75% | 6.16(1.55) | 5.29(1.93) | 5.29(2.27) |
| Fear | 12.37% | 6.47(1.82) | 5.77(2.09) | 5.82(2.18) |
| Neutral | 28.63% | 6.16(1.69) | 4.50(2.07) | 4.65(2.20) |

To assess subjective feelings of emotion related to black-and-white drawings of human faces, we recruited an independent sample of 400 participants (157 males; mean age ± SD = 22.06 ± 3.46 yrs) in an online survey. Sixteen black-and-white drawings of human male faces with neutral expressions were selected from web photo galleries. All images were transformed to 300 x 360 pixels on a grey background (RGB:122, 122, 122). Five subsets of faces with happy, sad, angry, disgust and fearful expressions were drawn based on each neutral face using Photoshop by an artist, as illustrated in Figure S2.

**Figure S2.**
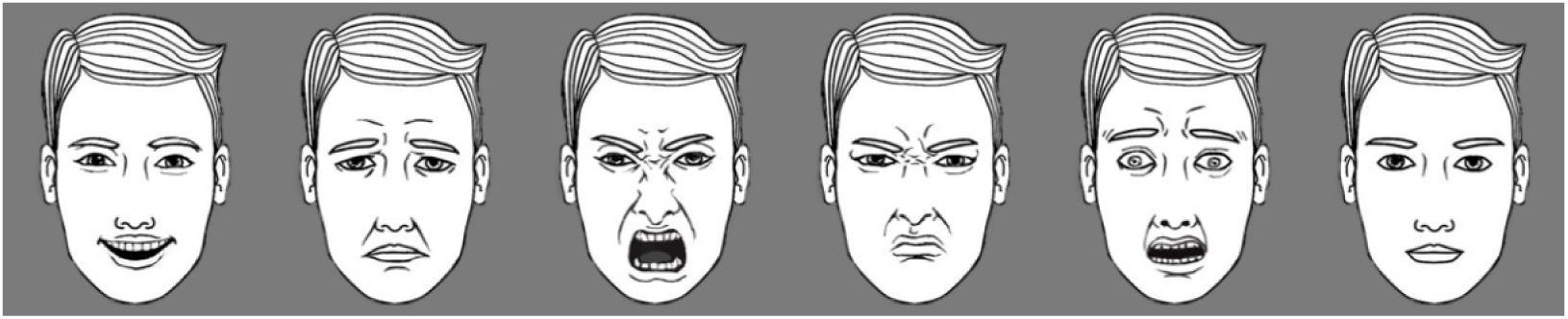
Illustrations of black-and-white drawings of human faces used in pilot Experiment 1.

Each participant was presented with faces of two models (each with 6 emotions) during the rating procedure similar to that used for drawings of skulls. The rating results, as shown in Table S2, suggest that the majority’s subjective feelings of emotion correspond to the facial expression of each set of black-and-white drawings of human faces. The emotional intensity and arousal related to these stimuli were slightly higher than moderate. Moreover, subjective ratings of emotion intensity and arousal were largely matched between skulls and emotional faces. We choose neutral and fearful faces as control stimuli in the following behavioral and brain imaging experiments because fearful faces were reported to induce the highest scores of death relatedness among the 5 types of facial expressions.

**Table S2.** Rating results related to drawings of faces in the pilot experiment

| Facial Expression | Frequency | Intensity<br>(Mean±SD) | Arousal<br>(Mean±SD) | Death Relatedness<br>(Mean±SD) |
| --- | --- | --- | --- | --- |
| Happy | 91.75% | 6.79(1.47) | 5.33(1.68) | 2.16(1.51) |
| Sad | 82.5% | 6.41(1.45) | 5.15(1.75) | 5.03(1.81) |
| Angry | 84.63% | 7.69(1.14) | 6.23(1.67) | 3.98(2.02) |
| Disgust | 68.38% | 6.83(1.28) | 5.59(1.44) | 3.60(1.82) |
| Fear | 85.63% | 7.19(1.42) | 5.86(1.92) | 5.86(1.77) |
| Neutral | 75.5% | 6.91(1.56) | 3.67(1.80) | 2.65(1.56) |

### Pilot Experiment 2: Semantic overlapping of word stimuli with *Death* and *Life*

In pilot Experiment 2, we recruited an independent sample of healthy adults (N = 36, 17 males, mean age ± SD = 20.94 ± 2.12 yrs) to test semantic overlapping of life-related words and word ‘Life’ and semantic overlapping of death-related words and word ‘Death’. Sixteen Chinese words of animal and plant names and 16 Chinese words related to death (each consists of two Chinese characters) with matching word frequencies were adopted from the previous work^46^ (see Table S3). Each trial began with the presentation of a fixation cross for 500 ms at the center of a screen. One of the three primes (i.e., word ‘Life’, ‘Death’, or a scrambled stimulus produced by combination of words ‘Life’ and ‘Death’) was then presented for 120 ms at the center of a screen. After a 50-ms interstimulus interval a word randomly chosen from those in Table S3 was presented for 200 ms at the center of the screen. The participants were asked to classify the word as being life-related or death-related by pressing one of two keys on a standard keyboard using the left or right index finger. There was an interval that randomly varied between 600-1000 ms after a button press before start of the next trial. The task instruction empathized both speed and accuracy. Each character subtended a visual angle of 1.8°×2.4° (width × height) at a viewing distance of 60 cm. There were 30 trials of each word category after each type of primes.

**Table S3.** Words used in pilot Experiment 2.

| Death-related words | English translations | Life-related words | English translations |
| --- | --- | --- | --- |
| 残骸 | debris | 兰花 | orchid |
| 毒气 | poison gas | 苍鹰 | hawk |
| 毒药 | poison | 草药 | herbs |
| 祭坛 | altar | 棕榈 | palm |
| 惨案 | tragedy | 狗熊 | bear |
| 忌日 | death anniversary | 大象 | elephant |
| 黄泉 | underworld | 枣树 | jujube tree |
| 阴间 | hell | 莲花 | lotus |
| 凶器 | murderous weapon | 青蛙 | frog |
| 阎王 | Hades | 芙蓉 | hibiscus |
| 杀手 | murderer | 杨柳 | willow |
| 坟场 | cemetery | 榆树 | elm |
| 癌症 | cancer | 狮子 | lion |
| 陵墓 | mausoleum | 毛驴 | donkey |
| 墓地 | graveyard | 红松 | Korean pine |
| 绞刑 | hanging | 乌龟 | turtle |

Response accuracies to categorize life-related and death-related words were high (>87.9% across all conditions) and did not differ significantly between the three priming conditions (F (2, 34) = 3.078, p = 0.059, η_p_^2^ = 0.153, 90%CI = [0,0.304]). To estimate the priming effects by controlling intrinsic difference in responses to life-related and death-related words, we calculated differential reaction times (ΔRTs) for each participant by subtracting RTs to life-related and death-related words preceded by the scrambled stimuli from those preceded by words ‘Life’ and ‘Death’. ΔRTs were then subject to a 2×2 repeated measures analysis (ANOVA) with Prime (Life vs. Death) and Word type (life-related vs. death-related) as within-subjects factors. The ANOVA of ΔRTs showed a significant 2-way interaction (F(1,35) = 5.261, p = 0.028, η_p_^2^ = 0.131, 90% CI = [0.008, 0.300]), indicating distinct patterns of priming effects pertaining to life-related words and death-related words. Post hoc comparison further confirmed significantly faster RTs to life-related words in ‘Life’ vs. scrambled priming condition (Mean difference = −12.62, p = 0.016, 95% CI = [−22.733, −2.506]) and significantly faster RTs to death-related words in ‘Death’ vs. scrambled priming condition (Mean difference = −10.91, p = 0.035, 95% CI = [−20.998, −0.821]). However, ‘Life’ vs. scrambled priming did not significantly influence RTs to death-related words (Mean difference = −0.59, p = 0.914, 95% CI = [−11.621, 10.438]) and ‘Death’ vs. scrambled priming did not significantly influence RTs to life-related words (Mean difference = −0.02, p = 0.997, 95% CI = [−12.335, 12.294], Figure S3). These results demonstrate semantic overlapping of life-related words with word ‘Life’ and semantic overlapping of death-related words with word ‘Death’.

**Figure S3.**
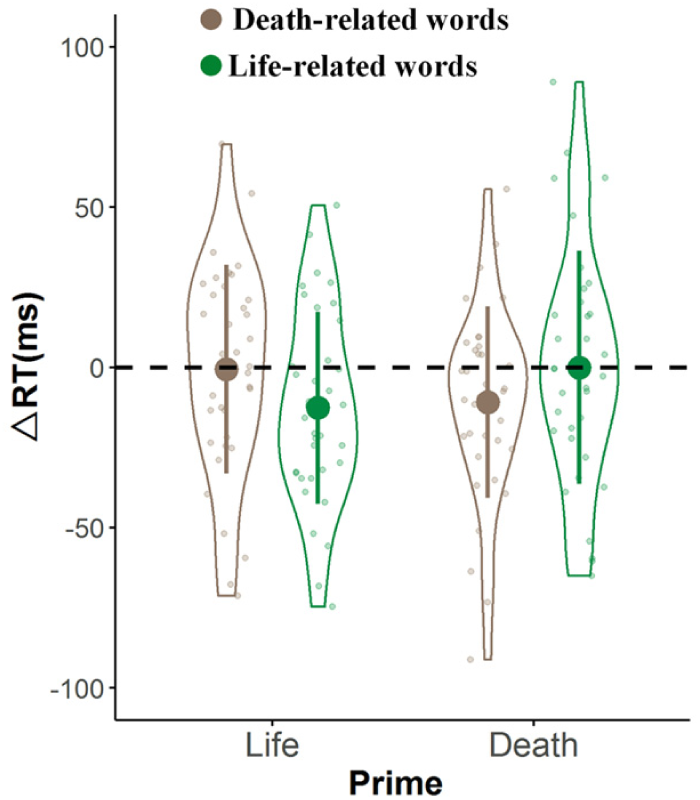
Results of priming effects on reaction times in pilot Experiment 2. Shown are group means (big dots), standard deviation (bars), measures of each individual (small dots), and distributions (violin shape) of ΔRTs.

### Experiment 1: Effects of perceived skulls on responses to life-related and death-related words

Thirty-eight university students (19 males, mean age ± SD = 20.87 ± 2.68 yrs) participated in Experiment 1 as paid volunteers. All participants self-reported no neurological diagnoses, nonreligious, right-handed and had normal or corrected-to-normal vision. Written informed consent was obtained prior to the experiment. This study was approved by the local ethics committee at the School of Psychological and Cognitive Sciences, Peking University. After the experiment, they were debriefed and explained the purpose of this study. A proper sample size was predetermined based on an assumed medium effect size (f = 0.25), and conventional power value (1-β = 0.95) and significant level (α = 0.05), using G*Power^47^, a sample size of 38 participants (with input parameters of correlation among repeated measurements = 0.48 and nonsphericity correlation = 1) was necessary to detect a significant two-way (2 × 4) interaction effect.

Prime stimuli consisted of 16 black-and-white drawings of skulls, 16 fearful faces, and 16 neutral faces that were evaluated in the pilot Experiment 1. We randomly choose one drawing from each type of the stimuli to create a scrambled stimulus. This procedure was repeated 16 times to obtain 16 scramble stimuli as control priming stimuli. Each trial began with the presentation of a fixation cross for 500 ms at the center of a screen. A prime, either a drawing of an upright or inverted skull, a fearful face, a neutral face, or a scrambled stimulus, was then shown for 120 ms at the center of a screen. After a 50-ms interstimulus interval a word randomly chosen from those in Table S3 was presented for 200 ms at the center of the screen. The participant was asked to classify the word as being life-related or death-related by pressing one of two keys on a standard keyboard using the left or right index finger within 1000 ms. There was an interval that randomly varied between 600-1000 ms after a button press before the next trial. The task instruction empathized both speed and accuracy. Each prime drawing subtended a visual angle of 2.4°×2.9° (width × height) and each character subtended a visual angle of 1.8°×2.4° (width × height) at a viewing distance of 60 cm. After performing 20 trials for practice, each participant completed 192 trials in 6 blocks of 40 trials (5 types of primes x 2 categories of words x 20 words in each category). The primes and words were presented in a random order. RTs and response accuracies were recorded in each condition.

Response accuracies were high (>93% in all conditions), as shown in Table S4. Thus, further data analyses focused on the results of RTs. Trials with incorrect responses or with RTs longer or shorter than mean RTs ± 3 SDs were excluded for further analysis. ΔRTs (i.e., RTs to words in the skull, neutral face, and fearful face conditions minus RTs in the scrambled stimulus condition) were subject to ANOVA with Prime (upright skull, inverted skull, fearful face, neutral face) and Word type (life-related vs. death-related) as within-subjects factors

**Table S4.**
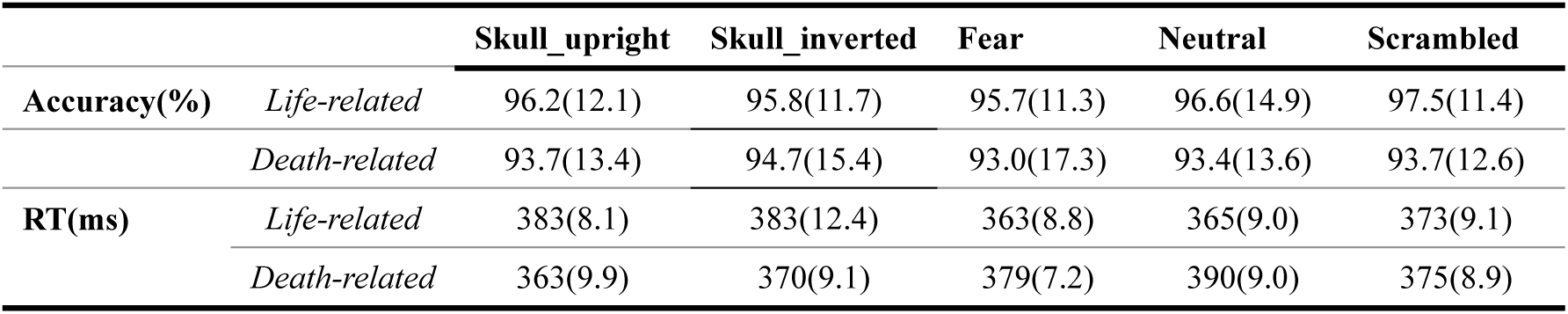
Response accuracy and reaction time (Mean (SE)) in Experiment 1

|  |  | Skull_upright | Skull_inverted | Fear | Neutral | Scrambled |
| --- | --- | --- | --- | --- | --- | --- |
| Accuracy(%) | *Life-related* | 96.2(12.1) | 95.8(11.7) | 95.7(11.3) | 96.6(14.9) | 97.5(11.4) |
|  | *Death-related* | 93.7(13.4) | 94.7(15.4) | 93.0(17.3) | 93.4(13.6) | 93.7(12.6) |
| RT(ms) | *Life-related* | 383(8.1) | 383(12.4) | 363(8.8) | 365(9.0) | 373(9.1) |
|  | *Death-related* | 363(9.9) | 370(9.1) | 379(7.2) | 390(9.0) | 375(8.9) |

### Replication of opposite skull priming effects on life- and death-related words in EEG/MEG experiments

We also collected RTs in the word classification task to test opposite skull priming effects on life-related and death-related words in the following EEG and MEG experiments (Experiments 3, 4, and 8).The stimuli and procedure were the same as those in Experiment 1 except that inverted skulls were excluded from the primes because the results in Experiment 1 showed that inverted skulls produced weak effects on RTs. ΔRTs were subject to a 3×2 ANOVA with Prime (skull, fearful face, neutral face) and Word type (life-related vs. death-related) as within-subject factors showed a significant interaction of Prime × Word type (F(2,170) = 26.23, p < 0.0001, η_p_^2^ = 0.236, 90% CI = [0.143,0.316]), post-hoc comparisons with Bonfferoni correction confirmed that skulls facilitated response death-related words but prolonged resposnes to life-related words (Mean difference = −30.41, p < 0.0001, 95% CI = [−17.35, −43.47]), whereas neutral faces showed reversed priming effects (Mean difference = 17.50, p = 0.008, 95% CI = [4.73, 30.26]). These results replicated the results in Experiment 1, providing further evidence for the opposite skull-priming effects on responses to life-related and death-related words.

**Figure S4.**
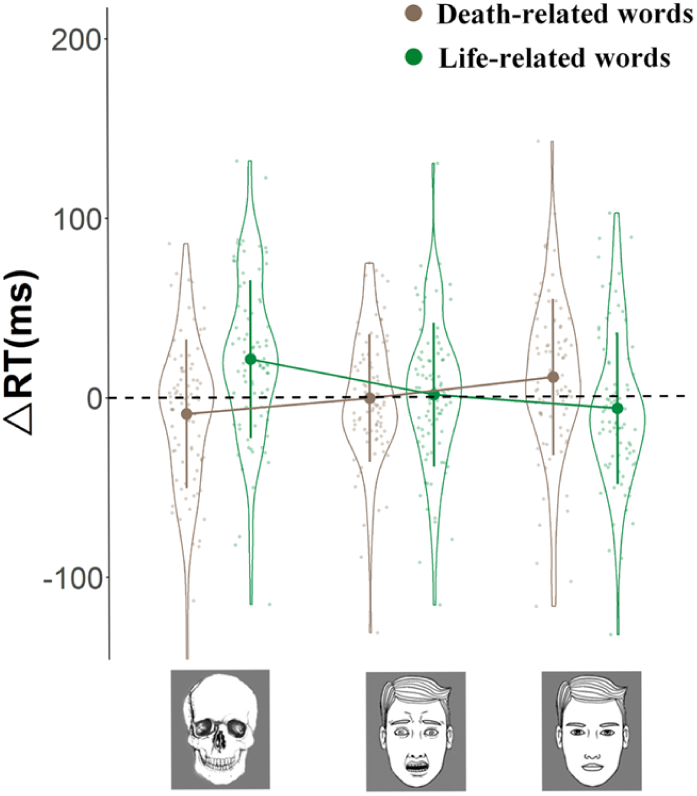
Behavioral results of priming effects across Experiments 3, 4, and 8. Shown are group means (big dots), standard deviation (bars), measures of each individual (small dots), and distributions (violin shape) of ΔRTs.

### Experiment 2: Effects of perceived houses on responses to life-related and death-related words

An independent sample (N = 38, 29 males, mean age ± SD = 20. 63 ± 2.02 yrs) was recruited in Experiment 2. The experimental design in Experiment 2 was the same as in Experiment 1 except that skulls were replaced by black-and-white drawings of houses (both Upright and Inverted). ΔRTs were subject to a 4×2 ANOVA with Prime (upright house, inverted house, fearful face, neutral face) and Word type (life-related vs. death-related) as within-subject factors showed a significant interaction of Prime x Word type (F (3,111) =3.01, p = 0.033, η_p_^2^ = 0.075, 90% CI = [0.003,0.144]). However, post-hoc comparisons failed to show any significant difference in ΔRTs to life-related and death-related words in each priming condition (House: Mean difference = 7.78, p = 0.473, 95% CI = [−13.95, 29.50]; Inverted house: Mean difference = −5.10, p = 0.673, 95% CI = [−29.40, 19.20]; Fearful faces: Mean difference = −9.03, p = 0.331, 95% CI = [−6.55, 34.52]; Neutral faces: Mean difference = 13.99, p = 0.176, 95% CI = [4.73, 30.26], Figure S5). The results suggest that inanimate objects do not necessarily produce opposite priming effects on life- and death-related words as skulls do.

**Figure S5.**
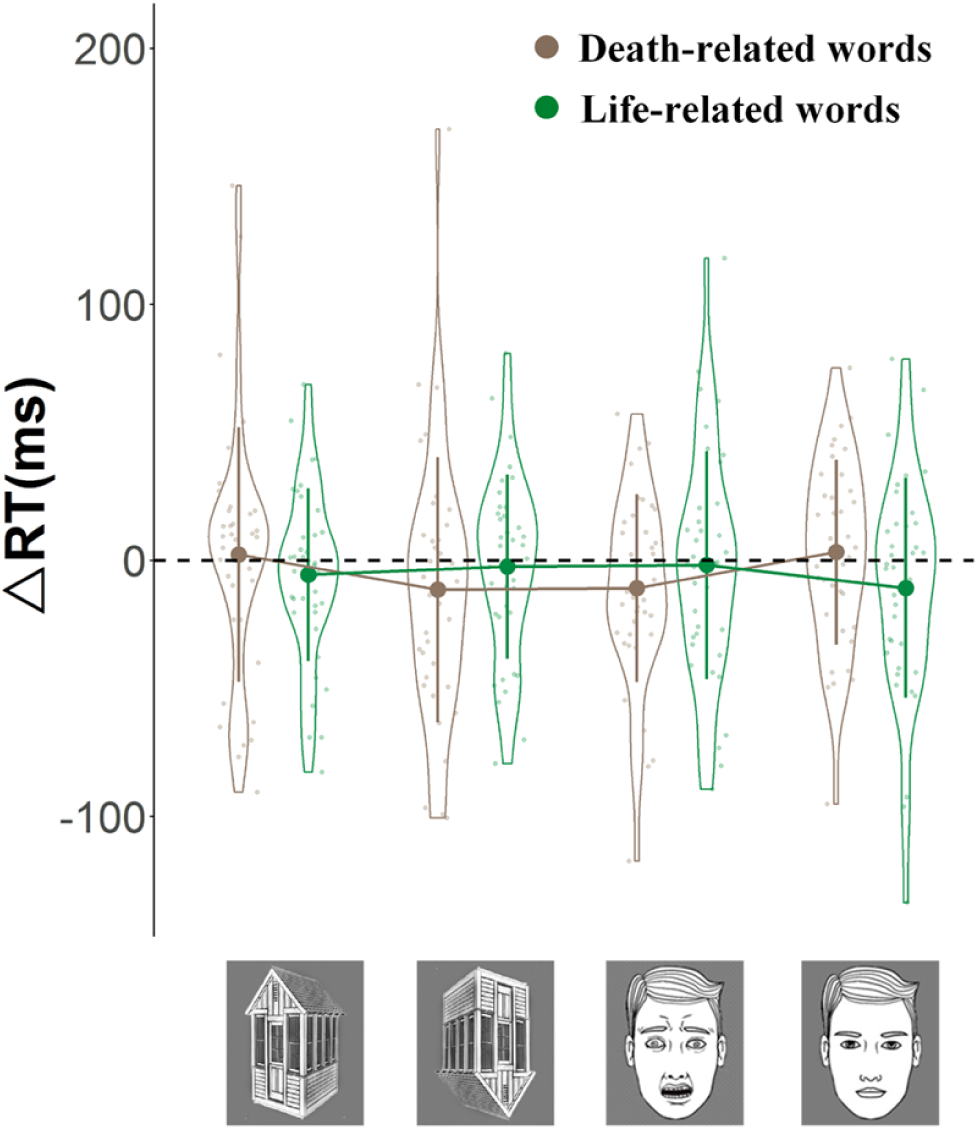
Behavioral results of priming effects of house and face stimuli in Experiment 2. Shown are group means (big dots), standard deviation (bars), measures of each individual (small dots), and distributions (violin shape) of ΔRTs.

### Experiment 3: Two-stage modulations of neural activity by perceived skulls Participants

Thirty university students (15 males; mean age ± SD = 20.93±2.00 yrs) participated in this study as paid volunteers. All participants self-reported no neurological diagnoses, unaffiliated with any religion right-handed and had normal or corrected-to-normal vision. Written informed consent was obtained prior to the experiment. The following brain imaging studies were approved by the local ethics committee at the School of Psychological and Cognitive Sciences, Peking University. After brain imaging data collection, all participants were debriefed by explaining the purpose, significance, and expected results of this study. Sample size was selected based on previous experiments from our lab and others using EEG/ERP.

### Stimuli and procedure

The participants were asked to complete two questionnaires prior to EEG recording given the functional role of cultural worldviews and self-esteem in buffering death anxiety. The Self-Construal Scale^48^ consists of 24 items for ratings on a 7-point Likert scale (1= strongly disagree, 7=strongly agree) for estimation of individuals’ cultural traits (i.e., independence and interdependence). The Self-Esteem Scale^49^ consists of 10 items for ratings on a 4-point Likert scale (1=strongly agree; 4=strongly disagree). However, these measures did not show reliable associations with neural responses to skulls and thus were not reported.

Participants were then asked to perform an one-back task on 16 black-and-white drawings of skulls, fearful faces, and 16 neutral faces, which were evaluated in the pilot Experiment 1. EEG was recorded continuously during the task. On each trial an image of a skull, a fearful face, a neutral face, or an inverted version of these stimuli was presented for 200 ms in the center of a gray background. It was followed by a fixation cross with a duration varying randomly from 800 to 1200 ms. Each stimulus subtended a visual angle of 2.4°×2.9° (width × height) at a viewing distance of 60 cm. The one-back task required participants to respond to the immediate repetition of the same stimulus in two successive trials by pressing a button. Each participant completed two EEG recording sessions. Upright and inverted skulls and neutral faces were shown in one EEG session (skull/neutral-face session). Upright and inverted fearful and neutral faces were shown in another EEG session (fearful/neutral-face session). There were 6 runs for each EEG session. Each run consisted of 84 trials (including 76 non-targets and 8 targets) with a 2-s break after every 12 trials. The order of the two sessions was counterbalanced across participants. There were 4 conditions in each EEG session (i.e., upright/inverted skulls and neutral faces, or upright/inverted fearful and neutral faces). Only non-target trials (114 trials in each condition) were included for EEG analysis.

After EEG recording, participants completed the Death Anxiety Scale (DAS)^50,51^ to assess individuals’ trait death anxiety. The DAS consists of 15 items rated using true or false classifications. The items were scored 0 or 1 and a higher score indicates a higher degree of death anxiety.

### EEG recording and data analysis

EEG was recorded from 64 Ag/AgCI scalp electrodes mounted in an elastic cap. The electrodes were arranged according to the international 10/20 system with an additional electrode below the right eye. EEG signals were collected with a band pass (0.1-100 Hz) and digitized with a sampling rate of 500 Hz (BrainAmp DC; Brain Products GmbH, Gilching, Germany), referenced online against FCz. AFz was used as the ground electrode. The impedance of each electrode was kept less than 5kΩ. EEG recorded at all electrodes was re-referenced to the average of the left and right electrodes (TP9/TP10) during off-line processing. After EEG was applied with a band-pass filter (0.1 – 40 Hz), epochs starting at 200 ms prior to the onset of a stimulus and lasting for 800 ms were extracted for non-target stimuli in each condition. The mean values from −200 to 0 ms served as the baseline interval for the respective epochs. These epochs were subjected to independent component analysis (ICA) using the EEGLAB (version 12) toolbox^52^ to detect and remove artifacts due to blinks and eye-movements. Subsequently, trials contaminated by residual artifacts exceeding ±70μV at any electrode were removed before further analyses. This procedure left 111.6±3.5 (mean ± SD) artifact-free trials per condition for further analyses. We then calculated ERPs to upright and inverted skulls and neutral faces separately in the skull/neutral-face session and to upright and inverted fearful and neutral faces in the fearful/neutral-face session, respectively. The mean amplitudes of each ERP component were calculated in the corresponding time window. The mean amplitude of death-related negativity (DRN) in response to perceived skulls was calculated at 120-140 ms at the occipital-parietal electrodes including O1, Oz, O2, P1, Pz, P2, PO3, POz, and PO4. The mean amplitude of the N170 components was calculated at 160-200 ms at the bilateral temporal-occipital electrodes including PO7, PO8, P7, and P8.

**Table S5.** Results of RTs and response accuracy (mean ± SD) in Experiment 3

| skull/neutral-face EEG session |  |  |  |  |
| --- | --- | --- | --- | --- |
|  | Skull |  | Neutral face |  |
|  | Inverted | Upright | Inverted | Upright |
| RT (ms) | 599± 69.9 | 554±79.2 | 602±70.7 | 585±67.8 |
| Accuracy(%) | 84.9±12.8 | 87.2±14.6 | 80.4±13.7 | 87.7±10.6 |
| fearful/neutral-face EEG session |  |  |  |  |
|  | Fearful face |  | Neutral face |  |
|  | Inverted | Upright | Inverted | Upright |
| RT (ms) | 574±69.4 | 595±67.9 | 573±66.8 | 569±101.4 |
| Accuracy(%) | 86±13.3 | 91±12.0 | 79.9±14.4 | 86.3±9.1 |

### ERP results

Perceived skulls and faces also elicited a positive wave at 100-120 ms over the bilateral occipitotemporal electrodes (P1). The P1 amplitudes were significantly enlarged by inverted than upright skulls and fearful faces (Main effect of Orientation and Stimuli F(1,29) = 6.59 and 17.34, p = 0.016 and p < 0.0001, η^2^_p_ = 0.185 and 0.374, 90% CI = [0.020, 0.372] and [0.141, 0.538]). However, the effect of stimulus inversion did not differ significantly between skulls and fearful faces (F(1,29) = 0.965, p = 0.334, η^2^_p_ = 0.032, 90% CI = [0.000, 0.180], Figure 2B). The P1 effect suggests enhanced perceptual processing of inverted than upright stimuli, which, however, was similar for skulls and fearful faces. The results of peak latencies, scalp distributions, and differential effects of the inversion manipulation on skulls and fearful faces indicate that the skull specific DRN cannot be simply explained as an extended P1 effect over the central occipto-parietal region.

We assessed whether the DRN amplitude varies across individuals’ self-esteem and cultural traits (i.e., self-construals) by conduction correlation analyses but did not find statistically significant results (DRN amplitude & scores of self esteem: r = 0.095, p = 0.625; DRN amplitude & scores of interdependence: r = −0.053, p = 0.787; DRN amplitude & scores of independence: r = 0.033, p = 0.866).

### Experiment 4: Replication of two-stage neural modulations by perceived skulls with controlled contextual stimuli

To compare neural response to perceived skulls and fearful faces in the same context, we recorded EEG from an independent sample (N = 28, 18 males; mean age = 20.43±1.68 yrs) in Experiment 4. The stimuli and procedure were the same as in Experiment 3 except that black-and-white drawings of upright and inverted skulls, fearful faces, and neutral faces were presented in the same block of trials. There were 7 blocks of 106 trials. RTs and response accuracies in the one-back ask are shown in Table S6.

**Table S6.** Results of RTs and response accuracy (mean ± SD) in Experiment 4

|  | skull |  | Fearful face |  | Neutral face |  |
| --- | --- | --- | --- | --- | --- | --- |
|  | Inverted | Upright | Inverted | Upright | Inverted | Upright |
| RT (ms) | 594±72.6 | 611±84.3 | 599±94.9 | 655±64.8 | 624±124 | 592±60.9 |
| Accuracy (%) | 84.6±12.6 | 91.2±11.4 | 85.7±13.3 | 92.7±10.7 | 80.1±17.1 | 89.6±11.4 |

Similar to those in Experiment 3, ERPs to upright (vs. inverted) skulls were characterized by a negative neural activity at 126-146 ms with the maximum amplitude over the middle occipitoparietal areas (Figure S6A). To quantify the DRN, we calculated the mean amplitude of the differential wave at 126-146 ms over the occipitoparieral electrodes in response to upright vs. inverted stimuli in each condition. The mean amplitudes of difference waves (upright minus inverted stimuli) were subject to ANOVAs with Stimuli (skull, fearful face, and neutral face) as within-subject variable. This revealed a significant main effect of Stimuli (F(2,54) = 32.82, p < 0.001, η^2^_p_ = 0.549, 90% CI = [0.380, 0.641]). Post-hoc comparisons with Bonfferoni correction showed that the DRN amplitude elicited by skulls was significantly larger than the mean amplitudes of difference waves in the DRN time window in response to fearful faces (Mean difference = −3.23 μV, p < 0.0001, 95% CI = [−4.71, −1.75]) and neutral faces (Mean difference = −3.18 μV, p < 0.0001, 95% CI = [−4.30, −2.06]). The mean amplitudes of difference waves, however, did not differ significantly between fearful and neutral faces (Mean difference = −0.045 μV, p = 0.31, 95% CI = [−0.75, 0.84]). These results replicated the DRN that differentiates between ERPs to skulls and fearful faces when these stimuli were presented in the same context of neutral faces. The results provide further evidence for an early modulations of brain activity by perceived skulls.

We also conducted an ANOVA of the mean amplitude of ERPs in the N170 time window (160-200 ms) with Stimuli (skull, fearful face, neutral face) and Orientation (upright vs. inverted) as within-subjects variables. There was a significant interaction of Stimuli x Orientation (F(2,54) = 14.37, p < 0.001, η^2^_p_ = 0.347, 90% CI = [0.165, 0.470]). Post-hoc comparisons with Bonfferoni correction showed that the N170 amplitude was significantly enlarged by upright than inverted skulls (Mean difference = −0.58 μV, p = 0.025, 95% CI = [−1.08, −0.08]), whereas the opposite effect was significant for the N170 amplitudes in response to fearful faces (Mean difference = 0.61 μV, p = 0.007, 95% CI = [0.18, 1.04]) and neutral faces (Mean difference = 0.79 μV, p < 0.0001, 95% CI = [0.41, 1.17], Figure S6B). These results replicated the findings of a specific pattern of modulation of the N170 amplitudes to perceived skulls and provide further evidence for a second-stage neural modulations by perceived skulls.

**Figure S6.**
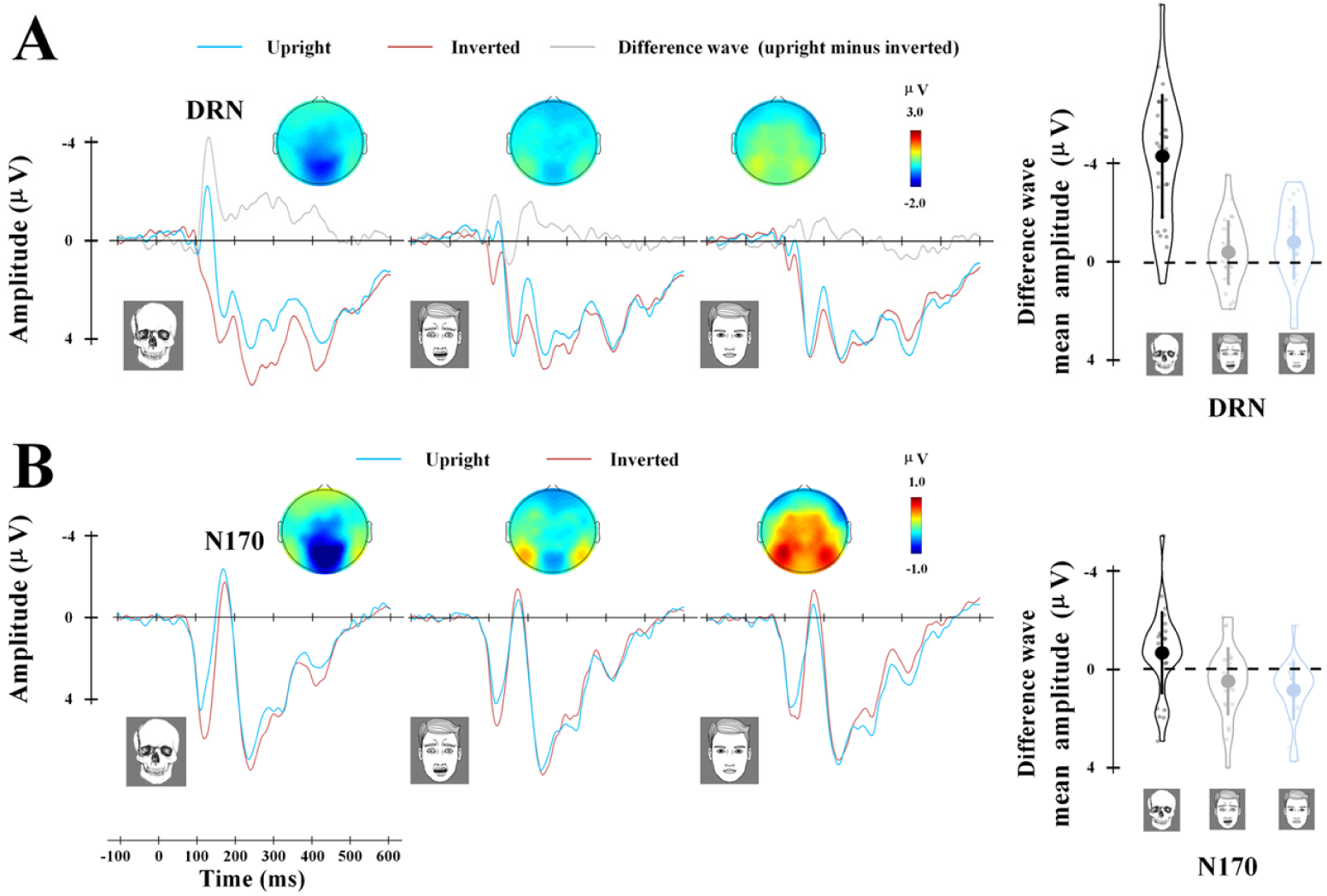
Results in Experiment 4. (A) ERPs to skull, fearful faces, and neutral faces at the occipito-parietal electrodes. (B) ERPs to skull, fearful faces, and neutral faces the lateral occipito-temporal electrode. The voltage topographies show scalp distributions of the difference wave (upright minus inverted skulls or fearful faces) in DRN and N170 time windows. The right panels illustrate the mean amplitudes of the differential wave in the DRN and N170 time windows (upright minus inverted). Shown are group means (big dots), standard deviation (bars), measures of each individual (small dots), and distributions (violin shape) of the mean amplitudes of the difference waves in the DRN and N170 time windows.

### Experiment 5: Two-stage neural modulations by perceived skulls but not inanimate objects

To examine whether perception of any inanimate objects produces neural modulations similar to those elicited by perceived skulls, in Experiment 5, we recorded EEG from an independent sample (N=36, three participants were excluded from data analyses due to low response accuracies or artifacts during EEG recording, leaving 33 participants for data analyses (15 males; mean age ± SD= 20.84 ± 2.05 yrs)). The stimuli and procedure were the same as in Experiment 4 except the following. A set of 16 black-and-white drawings of houses were selected from web photo galleries. There were 7 blocks of 122 trials during EEG recording. In each block, each set of stimuli (upright skulls, inverted skulls, upright neutral faces, inverted neutral faces, upright fearful faces, inverted fearful faces, and upright houses) were presented once. There were 10 targets in each block. RTs and response accuracies in each condition are shown in Table S7.

**Table S7.** Results of RTs and response accuracy (mean ± SD) in Experiment 5

|  |  | Neutral face |  | Fearful face |  | Skull |  |
| --- | --- | --- | --- | --- | --- | --- | --- |
|  | House | Inverted | Upright | Inverted | Upright | Inverted | Upright |
| RT(ms) | 619±102 | 634±92.9 | 612±95.9 | 631±106 | 613±93.1 | 641±105 | 608±99.8 |
| Accuracy(%) | 92.5±10.6 | 88.6±12.2 | 95.0±7.7 | 88.6±12.6 | 92.9±7.8 | 91.2±11.7 | 94.0±10.7 |

Similarly, ERPs to upright skulls were characterized by a negative activity at 122-142 ms over the middle occipitoparietal area (i.e., DRN, Figure S7A). We compared the ERP amplitudes in this time window in response to upright stimuli by conducting a one-way ANOVA with Stimuli (skull, fearful face, neutral face, and house) as a within-subjects variable, which showed a significant main effect of Stimuli (F(3,96) = 29.76, p < 0.0001, η^2^_p_ = 0.482, 90% CI = [0.348, 0.562]). Post-hoc comparisons further confirmed the more negative amplitude in this time window to upright skulls compared to other stimuli (skulls vs. fearful faces: MD = −1.99 μV, p < 0.0001, 95% CI = [−2.68, −1.31]; skulls vs. neutral faces: MD = −2.08 μV, p < 0.0001, 95% CI = [−2.94, −1.22]; skulls vs. houses: MD = −2.39 μV, p < 0.0001, 95% CI = [−3.32, −1.47], all Bonfferoni corrected). However, comparisons of the ERP amplitudes to upright faces and houses in the DRN time window did show any significant effect (houses vs. neutral faces, MD = 0.31 μV, p = 1.000, 95% CI = [−0.59, 1.22]; houses vs. fearful faces, MD = 0.40 μV, p = 0.886, 95% CI = [−0.36, 1.15]; neutral faces vs. fearful faces, MD = 0.08 μV, p = 1.000, 95% CI = [−0.50, 0.67]). These results indicate that the DRN was elicited by skulls but not by houses. We also conducted a one-way ANOVA of the mean amplitudes of the difference wave (upright minus inverted) with Stimuli (skull, fearful face, and neutral face) as a within-subjects variable, which similarly showed a significant main effect of Stimuli (F(2,64) = 76.99, p < 0.0001, η^2^_p_ = 0.706, 90% CI = [0.593, 0.765]). Post-hoc comparisons further confirmed that the skull-elicited DRN was significantly larger than the amplitudes of the difference wave elicited by neutral faces (MD = −2.69 μV, p < 0.0001, 95% CI = [−3.44, −1.95]) and by fearful faces (MD = −3.67, p < 0.0001, 95% CI = [−4.63, −2.72]).

**Figure S7.**
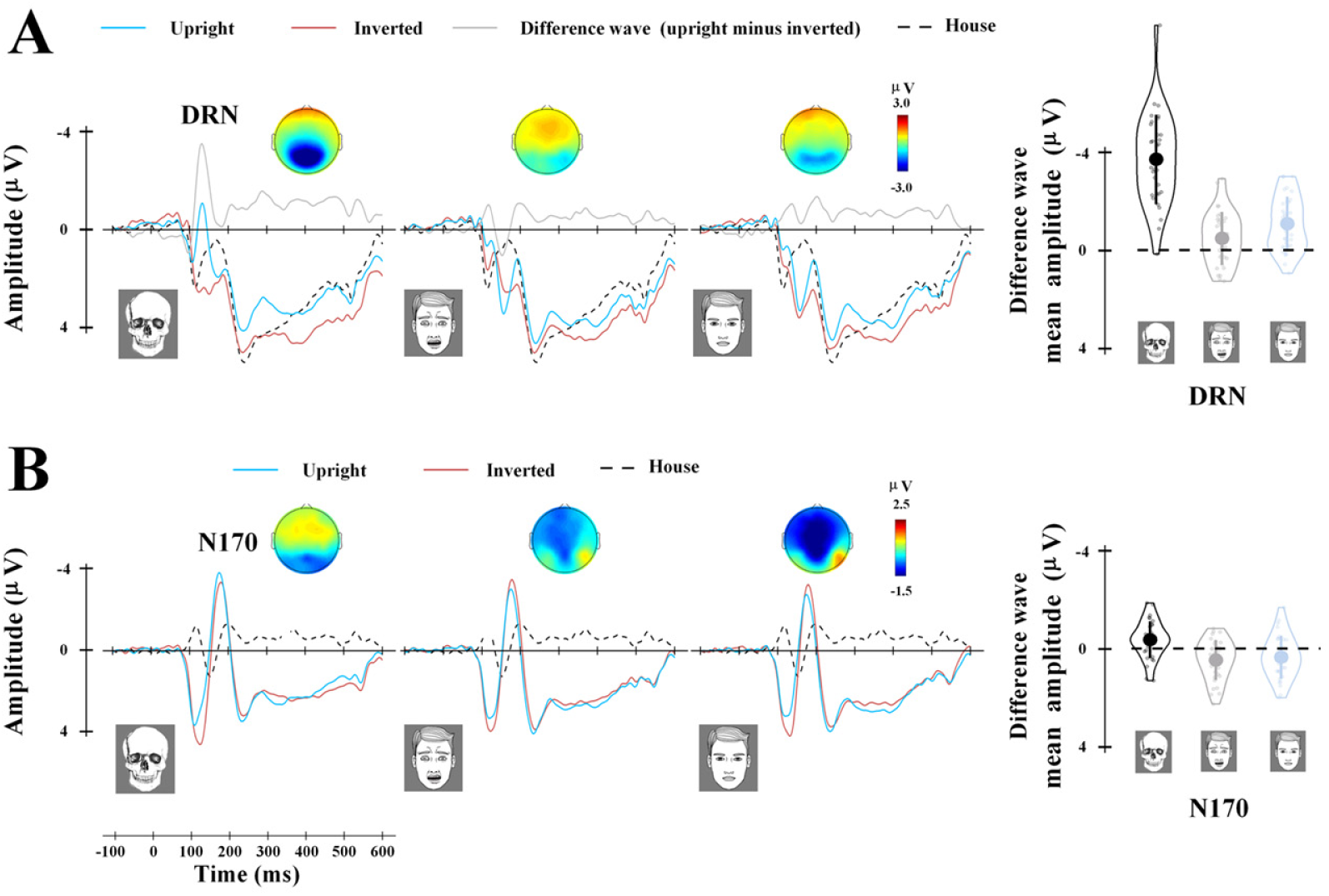
Results in Experiment 5. (A) ERPs to skull, fearful faces, neutral faces, and houses at the occipito-parietal electrodes. (B) ERPs to skull, fearful faces, neutral faces, and houses at the lateral occipito-temporal electrode. The voltage topographies show scalp distributions of the difference wave (upright minus inverted skulls or fearful faces) in DRN and N170 time windows. The right panels illustrate the mean amplitudes of the differential wave in the DRN and N170 time windows (upright minus inverted). Shown are group means (big dots), standard deviation (bars), measures of each individual (small dots), and distributions (violin shape) of the mean amplitudes of the difference waves in the DRN and N170 time windows.

We conducted a one-way ANOVA of the mean amplitudes in the N170 time window to upright stimuli with Stimuli (skull, fearful face, and neutral face) as a within-subjects variable. This showed a significant main effect of stimuli (F(3,96) = 35.50, p < 0.0001, η^2^_p_ = 0.526, 90% CI = [0.398, 0.601]). Post-hoc comparisons indicated that houses elicited a significantly smaller N170 amplitude compared to neutral faces (MD = 1.51, p < 0.001, 95% CI = [0.61, 2.42]), fearful faces (MD = 1.71, p < 0.0001, 95% CI = [0.76, 2.66]), and skulls (MD = 2.49, p < 0.0001, 95% CI = [1.65, 3.32], all Bonfferoni corrected, Figure S7B). We also conducted two-way ANOVA with Stimuli (skull, fearful face, neutral face) and Orientations (upright vs. inverted) as within-subject variables, which showed a significant interaction of Stimuli x Orientations (F(2,64) = 14.22, p < 0.0001, η^2^_p_ = 0.308, 90% CI = [0.145, 0.426]). Post-hoc comparisons further showed that the N170 amplitude was significantly enlarged by inverted than upright faces (neutral faces: MD = 0.43 p = 0.009, 95% CI = [0.12, 0.74]; fearful faces: MD = 0.53, p = 0.001, 95% CI = [0.24, 0.82]), whereas opposite effect was significant for the N170 amplitudes in response to skulls (MD = −0.30, p = 0.031, 95% CI = [−0.57, −0.03], all Bonfferoni corrected). Together, these results indicate that ERPs to skulls are characterized by the DRN and N170 enhancement effects. These results provide further evidence for the two-stage neural modulations by skulls.

### Experiment 6: Two-stage neural modulations by images of human skulls

To verify skull-specific DRN and N170 effects during perception of images of real human skulls and faces, in Experiment 6, we recorded EEG from an independent sample (N=30, 15 males; mean age ± SD = 21.50 ± 2.49 yrs) who performed the one-back task on images of real human faces and skulls.

The stimuli consisted of 16 images of human skulls selected from web photo galleries and 16 neutral male faces from Chicago Face Database^53^ and MR2 database^54^. The size and luminance levels were matched for images of skulls and faces. The experimental procedure and data acquisition and analyses were the same as those in Experiment 3 except the following. There were 4 blocks of 128 trials. In each block there were 32 trials showing upright skulls, inverted skulls, upright faces, or inverted faces. RTs and response accuracies in each condition are shown in Table S8.

**Table S8.** Results of RTs and response accuracy (mean ± SD) in Experiment 6

|  | skull |  | Neutral face |  |
| --- | --- | --- | --- | --- |
|  | Inverted | Upright | Inverted | Upright |
| RT (ms) | 550±90.1 | 566±71.0 | 582±173 | 539±73.3 |
| Accuracy(%) | 73.1±22.7 | 85.5±14.9 | 82.3±17.5 | 89.7±14.4 |

The ERP results showed that upright compared to inverted images of human skulls elicited a negative activity at 132-152 ms after stimulus onset with the maximum amplitude at the occipito-parietal region (F(1,29) = 76.948, p < 0.001, η^2^_p_ = 0.726, 90% CI = [0.555, 0.803]). This effect significantly distinguished between images of skulls and faces (F (1,29) = 42.081, p < 0.001, η^2^_p_ = 0.592, 90% CI = [0.372, 0.705]). The N170 amplitude at 160-200 ms in response to images of faces showed a significant inversion effect (F(1,29) = 8.477, p = 0.007, η^2^_p_ = 0.226, 90% CI = [0.039, 0.411]), being larger to inverted compared to upright faces. The N170 amplitude to skull, however, showed a significantly opposite effect (F(1,29) = 13.552, p = 0.001, η^2^_p_ = 0.318, 90% CI = [0.097, 0.492]), being larger to upright than inverted images of skulls. The distinct pattern of the N170 amplitudes was confirmed by a significant 2-way interaction of Stimuli (skull vs. face) x Orientation (upright vs. inverted stimuli) (F(1,29) = 17.804, p < 0.001, η^2^_p_ = 0.380, 90% CI = [0.147, 0.543]). These results replicate the findings reported in Experiments 3-5 and demonstrate that the two-stage neural modulations by skulls are independent of low-level perceptual features of stimuli.

### Experiment 7: Configural information of skulls and DRN

To investigate what image features of skulls are pivotal for the two-stage neural modulations, in Experiment 7, we asked the participants in Experiment 4 to perform the one-back task while viewing black-and-white drawings of skulls with eyeballs and neutral faces without eyeballs (Figure 3B). The stimuli consisted 16 black-and-white drawings of skulls with eyeballs and 16 neutral faces without eyeballs, which were modified from those used in Experiment 3. There were 7 blocks of 64 trials (16 upright skulls, 16 inverted skulls, 16 upright neutral faces, 16 inverted neutral faces) during EEG recording. There were 111.9±0.21 (mean ± SD) artifact-free trials per condition for further data analyses.

**Table S9.** Results of RTs and response accuracy (mean ± SD) in Experiment 7

|  | skull |  | Neutral face |  |
| --- | --- | --- | --- | --- |
|  | Inverted | Upright | Inverted | Upright |
| RT (ms) | 596±124 | 572±66.2 | 563±82.1 | 567±69.0 |
| Accuracy(%) | 76.0±17.1 | 87.8±11.9 | 81.9±14.2 | 88.4±11.4 |

### Experiment 8: Sources of dynamic neural responses to skulls

To examine the neural circuit involved in skull-induced DA and how its dynamic responses relate to behavioral priming effects on responses to skulls, we recruited 30 university students (15 males; mean age = 21.76±2.10 yrs) as paid volunteers in Experiment 8. All participants completed both MEG and fMRI sessions. One participant dropped out of this experiment due to personal reasons. Thus 29 participants were left in MEG/fMRI data analyses.

The stimuli and procedure in the MEG session were the same as those in Experiment 3. After MEG recording the participants completed the behavioral priming task as in Experiment 1. To localize the occipital face area (OFA), fusiform face area (FFA), and parahippocampal place area (PPA) in each individual participant, the participants were scanned using fMRI during a face perception task. The black-and-white drawings of 16 neutral faces and 16 houses used for passive viewing were the same as those in Experiment 5. We adopted a box-car design that consisted of 8 blocks of trials. Each block lasted for 28 s during which either neutral faces or houses were presented. On each trial a face (or house) was presented for 200 ms at the center of a gray screen followed by a white fixation cross with a duration of 350 ms. The order of face/house blocks were counterbalanced across participants. Two successive blocks were separated by a 9-s break. While viewing faces or houses participants were asked to focus on the center of screen and to press a button upon an occasional red fixation.

### fMRI/MRI data acquisition and analysis

Functional and anatomical image acquisition was conducted on a Siemens Trio 3.0 T MR scanner with a 12-channel phase-array head coil at the Center for MRI Research, Peking University. Multiband functional images were acquired by using T2-weighted, gradient-echo, echo-planar imaging sequences sensitive to BOLD contrast (matrix = 112×112, 62 slices, 2mm slice thickness, 2×2×2 mm^3^ voxel size; repetition time (TR) = 2000 ms, echo time (TE) = 30 ms, field of view (FOV) =22.4 × 22.4 cm, flip angle (FA) = 90°, scanning order: Interleaved, multiband accelerate factor = 2). A high-resolution anatomical T1-weighted image was acquired for each participant (256×256mm matrix, 192 slices, 1×1×1.00 mm^3^ voxel size; TR = 2530ms, TE=2.98ms, inversion time (TI)=1100ms, FOV = 25.6×25.6 cm, FA = 7°, scanning order: interleaved). Padded clamps were used to minimize head motion and earplugs were used to attenuate scanner noises.

Functional images were preprocessed using SPM12 software (the Wellcome Trust Centre for Neuroimaging, London, UK, http://www.fil.ion.ucl.ac.uk/spm). The functional images were first corrected for within-scan acquisition time difference slices, then realigned to the first scan. Six movement parameters (translation; x, y, z and rotation; pitch, roll, yaw) were obtained after motion correction. Then, all images were spatially normalized to the Montreal Neurological Institute (MNI) template, resampled to 3×3×3 mm^3^ voxels and spatially smoothed using an isotropic Caussian kernel with 8mm full-width half-maximum. To identify the brain regions sensitive to face stimuli, a fixed effect model was applied to each subject’s data by contrasting face stimuli with house at a threshold of P < 0.001 (voxel level, uncorrected). The reverse contrast was used to identify brain regions sensitive to houses. Consistent with the right hemispheric dominance of face processing, we identified the right FFA and OFA in 28 of 29 participants. Face-related voxels located in the lateral temporal portion of the fusiform gyrus were designated as the FFA (mean (SD) coordinates of peak voxel: x = 42 (3.1), y = −54 (5.6), z = −19 (2.6)). Face-related voxels located on the lateral surface of the inferior occipital gyrus were designated as the OFA (x = 43 (5.0), y =-75 (5.7), z = −12 (4.0)). House-related voxels located in the medial region of the ventral temporal cortex were designated as the PPA (x = 30 (2.2), y = −48 (7.1), z = −11 (3.6)). The regions of interest (ROIs) of each participant were defined as spheres with 5-mm-radius centered at the peak voxels of these regions. In order to identify ROIs on brain surface, the volume-based fMRI activity was projected to Freesurfer’s pial surface of the cortex using algorithms of the FSL toolbox resulting in the ROIs being defined in MEG source space. Segmentation of T1 image was conducted with automated algorithms provided in the FreeSurfer software package^55^ (http://surfer.nmr.mgh.harvard.edu/). The reconstructed anatomy was used in the forward model of the MEG analysis and as spatial reference for all co-registrations between fMRI and MEG.

### MEG data acquisition and analysis

#### Preprocessing

Cortical neuromagnetic activity was recorded using a whole-head MEG system with 102 magnetometers and 204 planar gradiometers (Elekta Neuromag TRIUX) in a magnetically shielded room. The MEG signals were sampled at 1 kHz with an online 0.1-330 Hz band-pass filter. To co-register the MEG data with MRI coordinates, 3 anatomical landmarks (nasion, left, and right pre-auricular points), 4 HPI coils and at least 200 points on the scalp were digitized using the Probe Position Identification system (Polhemus, VT, USA). During the MEG scan, the head coil positions were repeatedly recorded by electromagnetic induction at the beginning and end of each block. A Maxfilter software (Elekta-Neuromag, Helsinki, Finland) with temporal signal space separation (tsss) was first used to remove the external interference to the raw MEG data. For each condition, the head positions of each pair of blocks were coregistered in reference to the position of the first block using Maxmove (a sub-component of Maxfilter software) before combining the MEG data. The offline analysis of MEG data was performed using Brainstorm package^56^ (Version: 20-Mar-2019). Continuous MEG data were lowpass filtered at 100Hz with a notch filter of 50Hz. Eye blink artifacts were attenuated with signal space projection (SSP) by visually inspecting and removing the corresponding SSP component. Epochs were extracted for non-target stimuli with each epoch starting 200 ms prior to the onset of the respective stimulus and lasting for 600 ms. The mean of MEG signals recorded before stimuli onset (−200 to 0 ms) served as a baseline for the respective epochs. Subsequently, trials exceeding 3500fT at any sensor were removed before further analyses, resulting inclusion of 105.1±13.1 (mean ± SD) trials in each condition.

#### Source reconstruction

For source reconstruction, cortical currents were estimated using a distributed model consisting of 15,002 current dipoles from the combined time series of magnetometer and gradiometer signals using a linear inverse estimator (weighted minimum-norm current estimate, signal-to-noise ratio of 3, depth weighting of 0.5) separately for each condition and for each participant in an overlapping-sphere head model. Dipole orientations were constrained to the individual MRIs. Noise covariance matrix was acquired from 2 min empty-room MEG recordings collected daily before the experiment. For each of the 15002 vertices, normalized source activations were obtained by standardizing values to pre-stimulus intervals (−200 to 0 ms) (subtracting the mean and dividing by standard deviation of the baseline) and computing the absolute value. For the group analysis, individual source-space data were projected to a standard brain model (Colin27, 15,002 vertices) and a 3-mm full-width half-maximum (FWHM) Gaussian kernel was applied for spatial smoothing. Whole-brain analyses of significant effect (Upright minus Inverted condition) in source-space used the cluster based permutation t test performed on 15,002 vertices of the cortical surface model during 100-200 ms (time window was defined based on previous EEG results, a predefined threshold p < 0.001, two-tailed, 1000 iterations). The significant clusters were defined using a cluster-level threshold p < 0.05 with corrections for multiple comparisons.

We extracted time courses of source-space MEG signals from the ROIs (OFA, FFA, PPA) independently defined in each participant using fMRI. We then calculated a point-to-point Pearson correlation between the amplitude of neural responses to upright skulls in each ROI and skull priming effects on RTs to classify life-related and death-related words (RTs in the skull priming condition minus that in the scrambled priming condition).

#### A linear support vector machine analysis

We conducted multivariate analyses of MEG sensor-space signals using a linear support vector machine method (SVMs; http://www.csie.ntu.edu.tw/~cjlin/libsvm/) for classification of upright vs. inverted skulls and upright vs. inverted fearful faces independently for each participant. Epochs of MEG sensor-space signals were extracted for non-target stimuli with each epoch starting 200 ms prior to stimulus onset and lasting for 600 ms. MEG data at each time point (−100 to 400 ms) were arranged as 306 dimensional measurement vectors. We randomly grouped 8 trials of upright or inverted skulls into sub-averaging samples to produce n samples for each condition and time point. The leave-one-out cross-validation approach treated n-1 samples from each condition as training data set and the two left-out samples (e.g., one for upright skulls and one for inverted skulls) for testing the classifier performance. This process was repeated 100 times with random reassignment of the data to training and testing sets, yielding an overall decoding accuracy of the classifier. We conducted one-sample t-test to compare the decoding accuracy (vs. 50%) at each time point across participants. Group-level decoding was further estimated using cluster-based permutation t-tests using a predefined threshold p < 0.001, two-tailed, 10,000 iterations, and a cluster-level threshold p < 0.05 with corrections for multiple comparisons.

## Results

**Table S10.** Results of RTs and response accuracy (mean ± SD) in Experiment 8

| skull/neutra-face EEG session |  |  |  |  |  |
| --- | --- | --- | --- | --- | --- |
|  |  | Skull |  | Neutral face |  |
|  |  | Inverted | Upright | Inverted | Upright |
| RT (ms) | | 592 ± 82.0 | 559 ± 66.2 | 588 ± 72.1 | 588 ± 68.1 |
| Accuracy(%) | | 83.1 ± 15.3 | 88.7 ± 13.0 | 78.9 ± 20.3 | 86.0 ± 16.0 |
| fearful/neutra-face EEG session |  |  |  |  |  |
|  |  | Fearful face |  | Neutral face |  |
|  |  | Inverted | Upright | Inverted | Upright |
| RT (ms) | | 614 ± 73.7 | 590 ± 52.0 | 599 ± 61.7 | 589 ± 63.7 |
| Accuracy(%) | | 81.3 ± 17.3 | 85.9 ± 13.2 | 72.4 ± 18.6 | 83.8 ± 16.2 |

**Figure S8.**
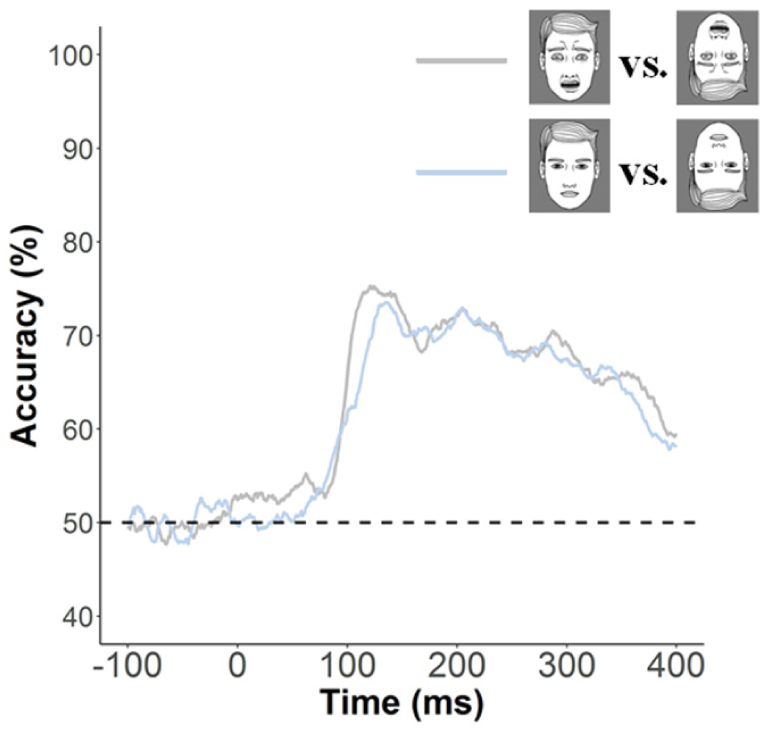
Results of decoding accuracies. We also calculated the accuracy of decoding upright vs. inverted neutral faces. The decoding accuracy was significantly higher than the chance level (50%) for upright vs. inverted faces at 80-400 ms (p < 0.001) in the skull/neutral face session and at 95-400ms (p < 0.001)in the fearful/neutral face session. Cluster-based t-tests confirmed a significantly higher decoding accuracy for upright vs. inverted skulls than upright vs. inverted neutral faces at 107-152 ms post-stimulus (p < 0.001), whereas similar analyses failed to show significant difference in the decoding accuracy between fearful and neutral faces.

**Figure S9.**
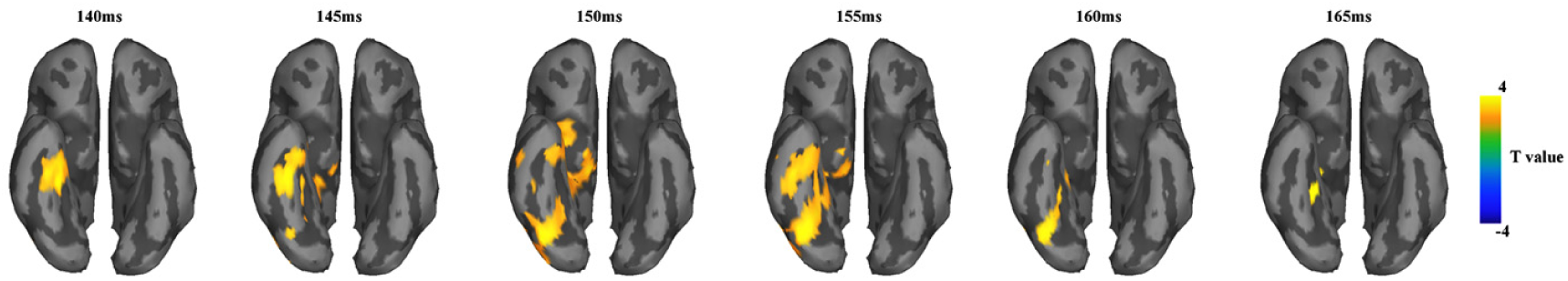
Illustrations of modulations of neural responses to upright vs. inverted neutral faces. Significant effects were identified using cluster-based permutation t-tests with a predefined threshold p < 0.001, two-tailed, 1000 iterations, and a cluster-level threshold p < 0.05 with corrections for multiple comparisons.

### Results of partial correlation analyses

To further confirm distinct patterns of the associations between OFA/FFA activity and skull-priming effects on behavioral responses to life-related and death-related words, we calculated partial correlations between mean OFA activity (105-150 ms) and ΔRTs to life-related words after controlling the variance of ΔRTs to death-related words and between mean FFA activity (166-188 ms) and ΔRTs to death-related words after controlling the variance of ΔRTs to life-related words. The results of these partial correlation analyses further confirmed distinct associations of the priming effects on life-related and death-related words with modulations of neural response to perceived skulls in two separate brain regions and two successive time windows (p< 0.01).

## References

1. Harris, P. L. Children’s understanding of death: from biology to religion. Philos Trans R Soc Lond B Biol Sci. 373, 20170266 (2018)..

2. Rank, O. Art and artist: Creative urge and personality and development. New York: Alfred A. Knopf (1932).

3. Pyszczynski, T., Kesebir, P. Culture, ideology, morality, and religion: Death changes everything. In P. R. Shaver & M. Mikulincer (Eds.), Meaning, mortality, and choice: The social psychology of existential concerns (p. 75–91) (2012).

4. Arndt, J., Vess, M., Cox, C. R., Goldenberg, J. L., Lagle, S. The psychosocial effect of thoughts of personal mortality on cardiac risk assessment. Med. Decision Making, 29, 175–181(2009).

5. Malpas, J. E., Solomon, R. C. Death and philosophy. Routledge (2002).

6. White, W., Handal, P. J. The relationship between death anxiety and mental health/distress. OMEGA-J. Death and Dying, 22, 13–24 (1991).

7. Jonas, E., Schimel, J., Greenberg, J., Pyszczynski, T. The Scrooge effect: Evidence that mortality salience increases prosocial attitudes and behavior. Pers. Soc. Psychol. Bullet. 28, 1342–1353 (2002).

8. Solomon, S., Greenberg, J., Schimel, J., Arndt, J., Pyszczynski, T. Human awareness of mortality and the evolution of culture. In The psychological foundations of culture. Psychology Press (pp. 24–49) (2003).

9. Arndt, J., Lieberman, J. D., Cook, A., Solomon, S. Terror Management in the Courtroom: Exploring the Effects of Mortality Salience on Legal Decision Making. Psychology, Public Policy, and Law, 11, 407–438 (2005).

10. Zaleskiewicz, T., Gasiorowska, A., Kesebir, P. Saving can save from death anxiety: Mortality salience and financial decision-making. PloS one, 8, e79407 (2013).

11. Luo, S., Shi, Z., Yang, X., Wang, X., Han, S. Reminders of mortality decrease midcingulate activity in response to others’ suffering. Soc. Cogn. Affect. Neurosci. 9, 477–486 (2014).

12. Luo, S., Wu, B., Fan, X., Zhu, Y., Wu, X., Han, S. Thoughts of death affect reward learning by modulating salience network activity. NeuroImage 202, 116068 (2019).

13. Silveira, S., Graupmann, V., Agthe, M., Gutyrchik, E., Blautzik, J., Demirçapa, I., HennigFast, K. Existential neuroscience: Effects of mortality salience on the neurocognitive processing of attractive opposite-sex faces. Soc. Cogn. Affect. Neurosci. 9, 1601–1607 (2014).

14. Li, X., Liu, Y., Luo, S., Wu, B., Wu, X., Han, S. Mortality salience enhances racial in-group bias in empathic neural responses to others’ suffering. NeuroImage 118, 376–385 (2015).

15. Gonçalves, A., Biro, D. Comparative thanatology, an integrative approach: exploring sensory/cognitive aspects of death recognition in vertebrates and invertebrates. Philos Trans R Soc Lond B Biol Sci. 373, 20170263 (2018).

16. Grant, A. M., Wade-Benzoni, K. A. The hot and cool of death awareness at work: Mortality cues, aging, and self-protective and prosocial motivations. Acad. Manag. Rev. 34, 600–622 (2009).

17. Han, S., Qin, J., Ma, Y. Neurocognitive processes of linguistic cues related to death. Neuropsychologia 48, 3436–3442 (2010).

18. Quirin, M., Loktyushin, A., Arndt, J., Küstermann, E., Lo, Y. Y., Kuhl, J., Eggert, L. Existential neuroscience: a functional magnetic resonance imaging investigation of neural responses to reminders of one’s mortality. Soc. Cogn. Affect. Neurosci. 7, 193–198 (2012).

19. Shi, Z., Han, S. Transient and sustained neural responses to death-related linguistic cues. Soc. Cogn. Affect. Neurosci. 8, 573–578 (2013).

20. Yanagisawa, K., Abe, N., Kashima, E. S., Nomura, M., Yanagisawa, K., Abe, N. Self esteem modulates amygdala-ventrolateral prefrontal cortex connectivity in response to mortality threats. J. Exp. Psychol. General 145, 273283 (2015).

21. Klackl, J., Jonas, E., Kronbichler, M. Existential neuroscience: self-esteem moderates neuronal responses to mortality-related stimuli. Soc. Cogn. Affect. Neurosci. 9, 1754–1761 (2014).

22. Kanjou, Y. Study of Neolithic human graves from Tell Qaramel in North Syria. Int. J. Mod. Anthrop. 2, 25–37 (2009)

23. Hallgren, F. Mesolithic skull depositions at Kanaljorden, Motala, sweden. Curr. Swed. Archaeol. 19, 244–246 (2011).

24. Barrett, H.C., Behne, T. Children’s understanding of death as the cessation of agency: a test using sleep versus death. Cognition 96, 93–108 (2005).

25. Kearl, M. C. The proliferation of skulls in popular culture: A case study of how the traditional symbol of mortality was rendered meaningless. Mortality 20, 1–18 (2015).

26. Taylor, C. C. Democritus and Lucretius on death and dying. In Democritus: *Science, the arts, and the care of the soul.* BRILL (pp. 77–86) (2007).

27. Grill-Spector, K., Weiner, K. S. The functional architecture of the ventral temporal cortex and its role in categorization. Nat. Rev. Neurosci. 15, 536–548 (2014).

28. Davison, A.C., Hinkley, D.V. Bootstrap methods and their application. UK: Cambridge University Press, Cambridge (1997).

29. Wyer, R. S., Jr., Sruli, T. K. Category accessibility: Some theoretical and empirical issues concerning the processing of social stimulus information. In E. T. Higgins, C. P. Herman, & M. P. Zanna (Eds.), Social cognition: The Ontario symposium. Hillsdale, NJ: Edbaum (Vol. 1, pp. 161–197). (1981).

30. Higgins, E. T., King, G. Accessibility of social constructs: Information-processing consequences of individual and contextual variability. In N. Cantor & J. E Kihlstrom (Eds.), Personality, cognition and social interaction. Hillsdale, N J: Erlbaum. (pp. 69–122) (2017).

31. Yovel, G. Neural and cognitive face-selective markers: an integrative review. Neuropsychologia 83, 5–13. (2016)

32. Balas, B., Koldewyn, K. Early visual ERP sensitivity to the species and animacy of faces. Neuropsychologia 51, 2876–2881 (2013).

33. Looser, C.E., Wheatley, T. The tipping point of animacy: How, when, and where we perceive life in a face. Psychol. Sci. 21, 1854–1862 (2010).

34. Baillet, S. Magnetoencephalography for brain electrophysiology and imaging. Nat. Neurosci. 20, 327–339 (2017).

35. Sha, L., Haxby, J. V., Abdi, H., Guntupalli, J. S., Oosterhof, N. N., Halchenko, Y. O., Connolly, A. C. The animacy continuum in the human ventral vision pathway. J. Cogn. Neurosci. 27, 665–678. (2015).

36. Proklova, D., Kaiser, D., Peelen, M. V. Disentangling representations of object shape and object category in human visual cortex: The animate–inanimate distinction. J. Cogn. Neurosci. 28, 680–692 (2016).

37. Gauthier, I., Tarr, M. J., Moylan, J., Skudlarski, P., Gore, J. C., Anderson, A. W. The fusiform “face area” is part of a network that processes faces at the individual level. J. Cogn. Neurosci. 12, 495–504 (2000).

38. Kanwisher, N., McDermott, J., Chun, M. M. The fusiform face area: a module in human extrastriate cortex specialized for face perception. J. Neurosci. 17, 4302-4311 (1997).

39. O’Craven, K. M., Downing, P. E., Kanwisher, N. fMRI evidence for objects as the units of attentional selection. Nature 401, 584–587 (1999).

40. Kanwisher, N., Yovel, G. The fusiform face area: a cortical region specialized for the perception of faces. Philos Trans R Soc Lond B Biol Sci. 361, 2109-2128. (2006).

41. Rossion, B., Hanseeuw, B., Dricot, L. Defining face perception areas in the human brain: a large-scale factorial fMRI face localizer analysis. Brain Cogn. 79, 138–157 (2012).

42. Zahn, R., Moll, J., Krueger, F., Huey, E. D., Garrido, G., Grafman, J. Social concepts are represented in the superior anterior temporal cortex. Proc. Natl. Acad. Sci. USA. 104, 6430–6435 (2007).

43. Zhou, Y., Gao, T., Zhang, T., Li, W., Wu, T., Han, X., Han, S. Neural dynamics of racial categorization predicts racial bias in face recognition and altruism. Nat. Hum. Behav. 4, 69–87 (2020).

44. Fan, X., Wang, F., Shao, H., Zhang, P., He, S. The bottom-up and top-down processing of faces in the human occipitotemporal cortex. eLife 9, e48764 (2020).

45. Anderson, J. R. Comparative thanatology. Curr. Bio. 26, R553–R556 (2016).

## References

46. Liu, X., Shi, Z., Ma, Y., Qin, J., & Han, S. (2013). Dynamic neural processing of linguistic cues related to death. PloS one, 8(6), e67905

47. Faul, F., Erdfelder, E., Buchner, A. & Lang, A. G. (2009). Statistical power analyses using G*Power 3.1: Tests for correlation and regression analyses. Behavioral Research Methods, 41, 1149–1160.

48. Singelis, T. M. (1994). The measurement of independent and interdependent self-construals. Personality and Social Psychology Bulletin, 20, 580–591.

49. Rosenberg, M. (1965). Society and the adolescent self-image, Princeton, NJ: Princeton University Press.

50. Templer, D. I. (1970). The construction and validation of a Death Anxiety Scale. Journal of General Psychology, 82, 165–177.

51. Lonetto, R., Fleming, S., & Mercer, G. W. (1979). The structure of death anxiety: a factor analytic study. Journal of Personality Assessment, 43, 388–392.

52. Delorme, A., & Makeig, S. (2004). EEGLAB: an open source toolbox for analysis of single-trial EEG dynamics including independent component analysis. Journal of Neuroscience Methods, 134, 9–21.

53. Ma, D. S., Correll, J., & Wittenbrink, B. (2015). The Chicago face database: A free stimulus set of faces and norming data. Behavioral Research Methods, 47, 1122–1135.

54. Strohminger, N., Gray, K., Chituc, V., Heffner, J., Schein, C., & Heagins, T. B. (2016). The MR2: A multi-racial, mega-resolution database of facial stimuli. Behavioral Research Methods, 48, 1197–1204.

55. Fischl, B., Van Der Kouwe, A., Destrieux, C., Halgren, E., Ségonne, F., Salat, D.H., Busa, E., Seidman, L. J., Goldstein, J., & Kennedy, D. (2004). Automatically parcellating the human cerebral cortex. Cerebral Cortex, 14, 11–22.

56. Tadel, F., Baillet, S., Mosher, J. C., Pantazis, D., & Leahy, R. M. (2011). Brainstorm: a user-friendly application for MEG/EEG analysis. Computational Intelligence and Neuroscience, 2011.

